# Distinct Connectivity Signatures of Emotions Enhance Precision of Network Biomarkers in Mood Disorders

**DOI:** 10.1101/2025.03.02.641022

**Authors:** Shuyue Xu, Linling Li, Ting Luo, Li Zhang, Benjamin Becker, Zhen Liang

## Abstract

Mood disorders, including Major Depressive (MDD) and Bipolar (BD) Disorder, are highly prevalent and debilitating conditions that contribute significantly to the global disease burden. These disorders are characterized by persistent emotional dysregulations, such as pervasive sadness and anhedonia, resulting in substantial functional impairments. Although neuroimaging studies have identified differences in brain activity and connectivity between individuals with MDD (MDDs) or BD (BDs) and healthy controls (HCs), reliable and reproducible neurofunctional markers for clinical diagnosis and treatment remain elusive. This study seeks to address this gap by introducing a novel approach that utilizes Divergent Emotional Functional Networks (DEFN), derived from functional magnetic resonance imaging (fMRI) during dynamic emotional processing in naturalistic contexts. Using a combination of naturalistic induction of sustained emotional experience with dynamic functional connectivity (dFC) and machine learning techniques, we decoded emotion-specific patterns of happiness and sadness in healthy individuals. Based on the dynamic functional connectivity signatures, we identify the DEFN and applied it to large clinical mood disorder datasets, including MDD (n=63) and BD patients (n=61). The model with DEFN demonstrated significant improvements in classification accuracy compared to conventional baseline models, achieving up to 10.75% and 9.92% performance increases in MDD and BD datasets, respectively. Additionally, DEFN were found to be highly reproducible across age, gender and models from emotion dataset, supporting the robustness of this model in distinguishing mood disorders from healthy controls. In conclusion, the DEFN approach presents a promising, reproducible, and clinically relevant neural marker for diagnosing and understanding emotional dysfunction in mood disorders, offering potential for more effective and timely interventions.

## Main

Mood disorders are highly prevalent and debilitating conditions which have become the leading contributor to the burden of disease collaborators (Collaborators, 2022). Major depressive (MDD) and bipolar disorder (BD) are among the most common mood disorders (Kessler, *et al*., 2003). These disorders are characterized by persistent emotional dysregulation, including pervasive sadness, or loss of interest or pleasure in activities once enjoyed (anhedonia) (Cromby and Willis, 2016) (Malhi, *et al*., 2021). Both lead to significant functional impairments and decreased quality of life (Kessler, *et al*., 2003, Toh, *et al*., 2015). Several studies have combined case-control neuroimaging designs or meta-analytic approaches to demonstrate differences in brain activity or connectivity between individuals with MDD or BD and healthy individuals (Li, *et al*., 2022). (Bore, *et al*., 2024) (Bore, *et al*., 2024) (Xu, *et al*., 2021). While these studies have provided valuable insights, they have failed to establish reliable and reproducible neurofunctional markers that can be used for clinical diagnosis or treatment selection (Etkin, 2019). While large scale studies have employed resting state fMRI and brain structural approaches to determine diagnostic markers for depression results have been highly inconsistent (Winter, *et al*., 2024). Limitations of this top-down approach using case-control studies have long been discussed and have for instance inspired the development of the Research Domain Criteria Approach (RDoC), a bottom-up approach with a focus on the underlying neurobehavioral domains (Insel, *et al*., 2010) (Etkin, 2019).

One major limitation that impedes bridging the behavioral domain approach with the prevailing focus on large scale resting state data is the lack of emotion processing domain-specific markers that can inform the detection of emotional dysfunction in mood disorders. Despite progress in the development of emotion-specific neural signatures in affective neuroimaging using task-based fMRI (Liu, *et al*., 2024) (Čeko, *et al*., 2022) the focus on experimental paradigms using sparse emotional stimuli and analytic procedures focusing on activation patterns has impeded translation into dynamic naturalistic contexts as well as development of disorder markers that capitalize on network level signatures of affective dysregulations that are at the core of the clinical symptoms of the disorders (Taschereau-Dumouchel, *et al*., 2022) (Zhou, *et al*., 2023). The few studies that explored emotion-specific network markers commonly focused on static connectivity (Saarimäki, *et al*., 2022) (Xu, *et al*., 2023) (Zhou, *et al*., 2023), which may not fully capture the dynamic, emotion-specific neural networks and may lack the replicability and robustness that have been demonstrated for dynamic connectivity measures (Abrol, *et al*., 2017). Despite progress in neuroimaging research, promising reports of clinically meaningful emotional-dysfunction marker in these disorders were not replicated, or only partially replicated (Pilmeyer, *et al*., 2022, Welton, *et al*., 2020). This results in prolonged and ineffective treatment, poorer prognoses, and increased healthcare costs. (Costello, *et al*., 2002). To address these limitations, we propose a novel two-step approach that fuses recent progress in the development of emotion-specific network markers under naturalistic emotion experience with network-based diagnosis of mood disorders based on dynamic resting state indices (Divergent Emotional Functional Networks, DEFN) in MDD and BD. By providing a more reproducible and clinically relevant neural marker, this approach has the potential to significantly enhance early-stage diagnostic accuracy, ultimately paving the way for more effective and timely interventions.

Functional magnetic resonance imaging (fMRI) is widely used in affective and psychiatric neuroscience to determine how the brain processes emotions and dysregulation in these processes in mental disorders (Gan, *et al*., 2022) (Liu, *et al*., 2024) (Čeko, *et al*., 2022). While paradigms using tasks to induced specific emotional states have been extensively utilized in healthy subjects, the vast majority of studies in patients has focused on resting-state fMRI (rs-fMRI) given the easier implementation into the clinical context and reduced burden for the patients (Canario, *et al*., 2021) (Parkes, *et al*., 2020) (Mousavian, *et al*., 2021). While some of the current limitations of rs-fMRI relate to technical issues inherent to the technique, e.g. head motion, wakefulness or replicability (Huskey, *et al*., 2018) (Wang, *et al*., 2017), rs-fMRI is limited dues to the lack of specificity for specific emotional states. Recent developments have therefore proposed novel naturalistic paradigms during which individuals are exposed to movies or narratives to approximate real-life environments and as such provide a more realistic reflection of brain activity during emotional experiences in ecological environments (Noah, *et al*., 2015) (Zhang, *et al*., 2022) (Xu, *et al*., 2023) (Sonkusare, *et al*., 2019). These naturalistic paradigms offer new insights into on how the human brain operates in real life, which is more dynamic and complex than the abstract tasks designed for laboratory settings. From a clinical perspective, the naturalistic paradigm share similar advantages with the resting-state paradigm in terms of participant compliance but imposes implicit behavioral constraints. Specifically, naturalistic paradigm fMRI significantly alleviates anxiety related to scanner performance and head movement, increases participant engagement and synchronization between subjects, allowing for more targeted research into mood disorders (Hasson, *et al*., 2004) (Eickhoff, *et al*., 2020).

Functional connectivity (FC) reflects the information integration and interaction between different brain regions and large-scale networks by observing the synchrony of blood-oxygen-level-dependent (BOLD) signals across these brain areas (Van Den Heuvel and Pol, 2010) (Bijsterbosch, *et al*., 2017). FC analysis usually assumes that brain activity remains stable throughout the data acquisition of several minutes while connectivity changes dynamically over the period. This static analysis may overlook subtle brain connectivity features that vary over time. Therefore, dynamic functional connectivity (dFC) has increasingly attracted interest in recent years, as it can detect transient brain functional connectivity patterns and reveal dynamic transitions between different brain states (Preti, *et al*., 2017) (Qiao, *et al*., 2020) (Jiang, *et al*., 2021). DFC - as an effective tool for exploring the integration of complex brain networks - has been widely applied in the research of mental neurological disorders, including mood disorders (Liu, *et al*., 2021) (Liu, *et al*., 2023) (Zhong, *et al*., 2022), schizophrenia (Du, *et al*., 2018) (Guo, *et al*., 2018), Attention Deficit Hyperactivity Disorder (ADHD) (Firouzi, *et al*., 2024), and Alzheimer’s Disease (AD) (Zhao, *et al*., 2022) (Matsui and Yamashita, 2023). For example, Lu et al. revealed that the disrupted dFC variability could distinguish patients with depressive episode from those with MDD with 83.44% classification accuracy (Lu, *et al*., 2023). Firouzi et al. have demonstrated that the model performance improved by up to 10% in classifying ADHD from typically developing children (TDC) using the dFC approach compared to the FC-based classification. Zhao et al. used dFC strength and a machine learning method to classify AD patients and healthy controls, finding differences in dFC strength in the left precuneus between the two groups (Zhao, *et al*., 2022).

We here capitalized on recent progress in naturalistic neuroimaging, dynamic network analyses and neurofunctional decoding across multiple datasets to (1) determine dynamic network-level signatures for emotional states in health individuals, and to next (2) translated these into clinical application to decode emotion-specific diagnostic markers based on rs-fMRI data acquired in large samples of individuals with BD and MDD. Happiness and sadness are core emotional dysfunctions in both MDD and BD, yet they manifest differently between the disorders (Fekadu, *et al*., 2017). MDD is primarily characterized by a persistent inability to experience positive emotions (anhedonia) and an exaggerated response to sadness (Pitsillou, *et al*., 2020), whereas BD involves dysregulated mood shifts, with heightened responses to both positive and negative emotions (Dodd, *et al*., 2019). Understanding these distinctions is critical for developing emotion-specific biomarkers, justifying our focus on happiness and sadness in this study. Our study was conducted in two stages. **(1) Stage 1: training the DEFN in healthy individuals.** A cohort of 52 healthy individuals underwent a naturalistic (movie) emotion induction paradigm using long movie clips designed to elicit happiness and sadness. Specifically, we curated a set of 12 different 10-minute movie clips (6 happiness and 6 sadness movies) and simultaneously recorded whole-brain fMRI responses (details see Xu et al., 2023). The dFC was assessed using a whole-brain region-of-interest (ROI) template comprising 400 cortical and 32 subcortical areas. Cross-subject and cross-episode emotion classification models were developed based on these dFC features, enabling the identification of the DEFN. DEFN is considered as a functional network signature representing emotion-specific brain dynamics in healthy individuals. **(2) Stage 2: translating DEFN to clinical application in mood disorder**. To assess the clinical utility of the DEFN, we applied it to rs-fMRI datasets from 286 individuals, including MDD dataset (n=174, MDDs=63, HCs=111) and BD dataset (n=112, BDs=61, HCs=51). Using leave-one-subject-out cross-validation and a linear support vector machine (SVM) model, we classified MDD/BD patients from healthy participants to validate the effectiveness of the DEFN. In the present work, we aimed to robustly identify the DEFN to uncover the emotion-specific dysfunctional patterns underlying MDD and BD symptomatology and in turn inform diagnostics and personalized interventions.

## Results

The entire analysis is illustrated in Fig 1. We employ sliding time window and k-means clustering methods to estimate four states that reflect dynamic brain activity during emotional episodes. For each episode, the average states are calculated, and their upper triangular matrix elements are extracted as one sample. To identify the DEFN, these samples are used to train an optimal linear SVM model that distinguishes between happiness and sadness in healthy individuals, along with its stable functional features that appear in each fold of cross-validation. The feature union derived from stable functional features appearing across states is used to calculate the network weights (i.e., DEFN) by calculating the ratio of the number of stable features to the total number of features within/between each network. Both a publicly available dataset and a self-acquired BD dataset, masked with DEFN information, are used to explore the emotion dysfunctional patterns in the two common mood disorders. Finally, the performance of the optimal model with DEFN is compared to the baseline model with whole brain information to assess the model efficacy, and the emotion dysfunctional patterns of MDD and BD are identified.

**Fig. 1:**
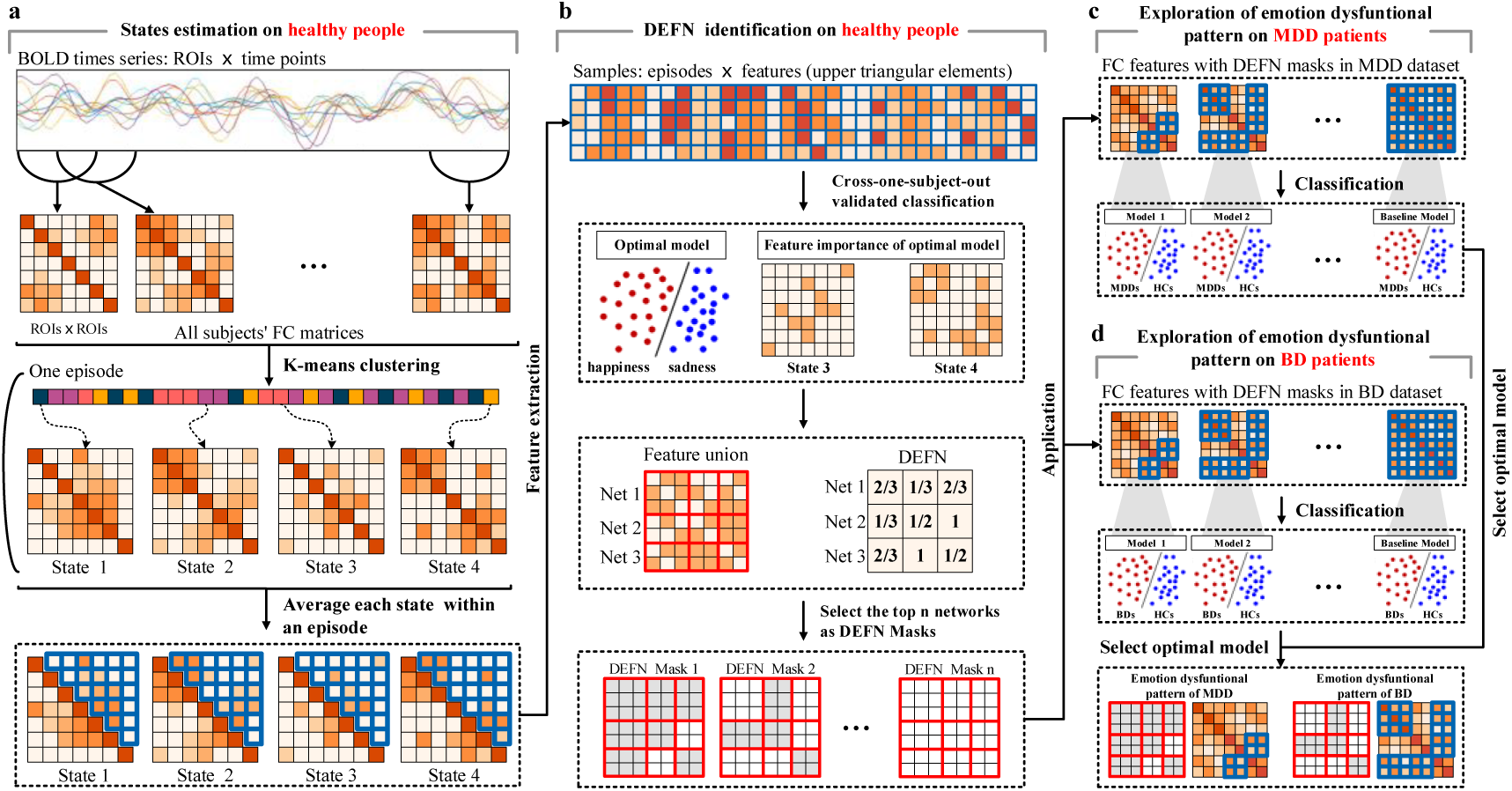
The flowchart of the entire analysis. a, Procedure of states estimation with the whole brain template of 432 ROIs on healthy individuals. b, DEFN Identification on healthy individuals in self-acquired emotion dataset (n=52). c, Exploration of emotion dysfunctional patterns of MDD in public MDD (n=174). d, Exploration of emotion dysfunctional patterns of BD in self-acquired BD datasets (n=112).

### Identified States and DEFN in Healthy Participants

The optimal number of clusters was estimated using the elbow method and determined resulting in four clusters. The four recurrent states reflecting brain function during long-term emotional experiences are depicted in Fig 2. Across all subjects and time points, the frequencies of states1 through 4 are approximately 5.87%, 19.81%, 35.46%, and 38.87%, respectively.

**Fig. 2:**
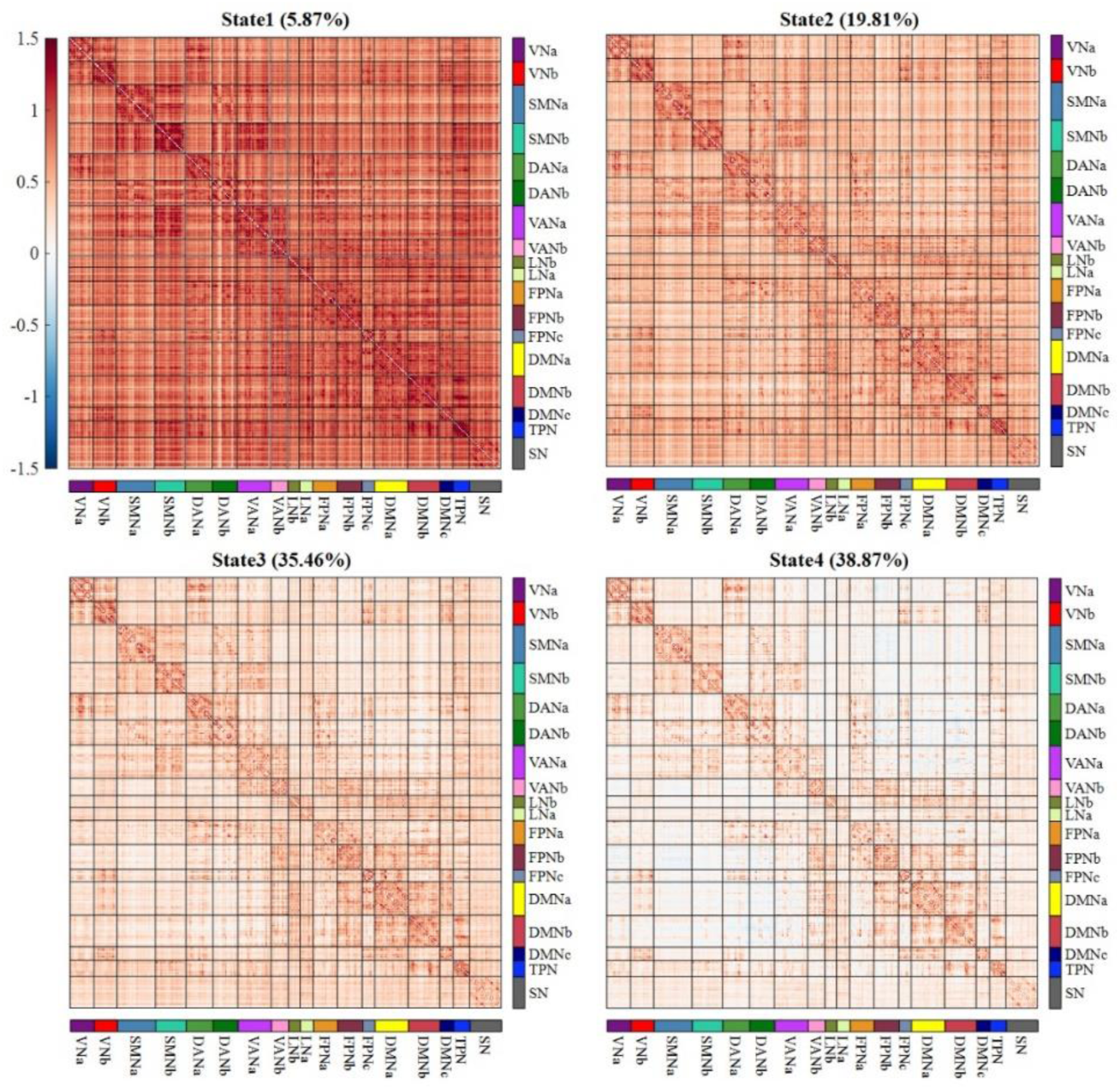
Results of state estimation on healthy participants. Centroids depicted the characteristics of four states, with the numbers in parentheses representing the proportions of these states across all participants’ time points.

Based on the obtained four states, the features were extracted by considering the connectivity strength within each state and integration of connectivity strengths across four states. A linear SVM model, combined with leave-one-subject-out cross-validation, was employed to decode emotional experiences of happiness and sadness successfully. To determine the optimal model, we identified the model with the highest classification accuracy and smallest standard deviation of accuracy as the optimal model. Based on the optimal model, the stable features that consistently emerged across per-fold cross-validation and the feature union of these stable features across all states were obtained. Finally, based on these stable features, the feature weights for each network were calculated.

The classification results (Table 1) show that the optimal model with best discrimination between happiness and sadness was built on combination features from state34 (accuracy: 83.99%). The statistical results indicate that the model performance based on states3, state4, state23, and state234 does not show a statistical difference compared to state34 (Defined as the second-best model). The present results indicate that in healthy individuals, happiness and sadness can be accurately discriminated using a linear SVM model with high accuracy of 83.99%. This high accuracy is primarily attributed to connections spanning between-network pairs associated with VNa and LNb (as shown in Fig.3 a).

**Fig. 3:**
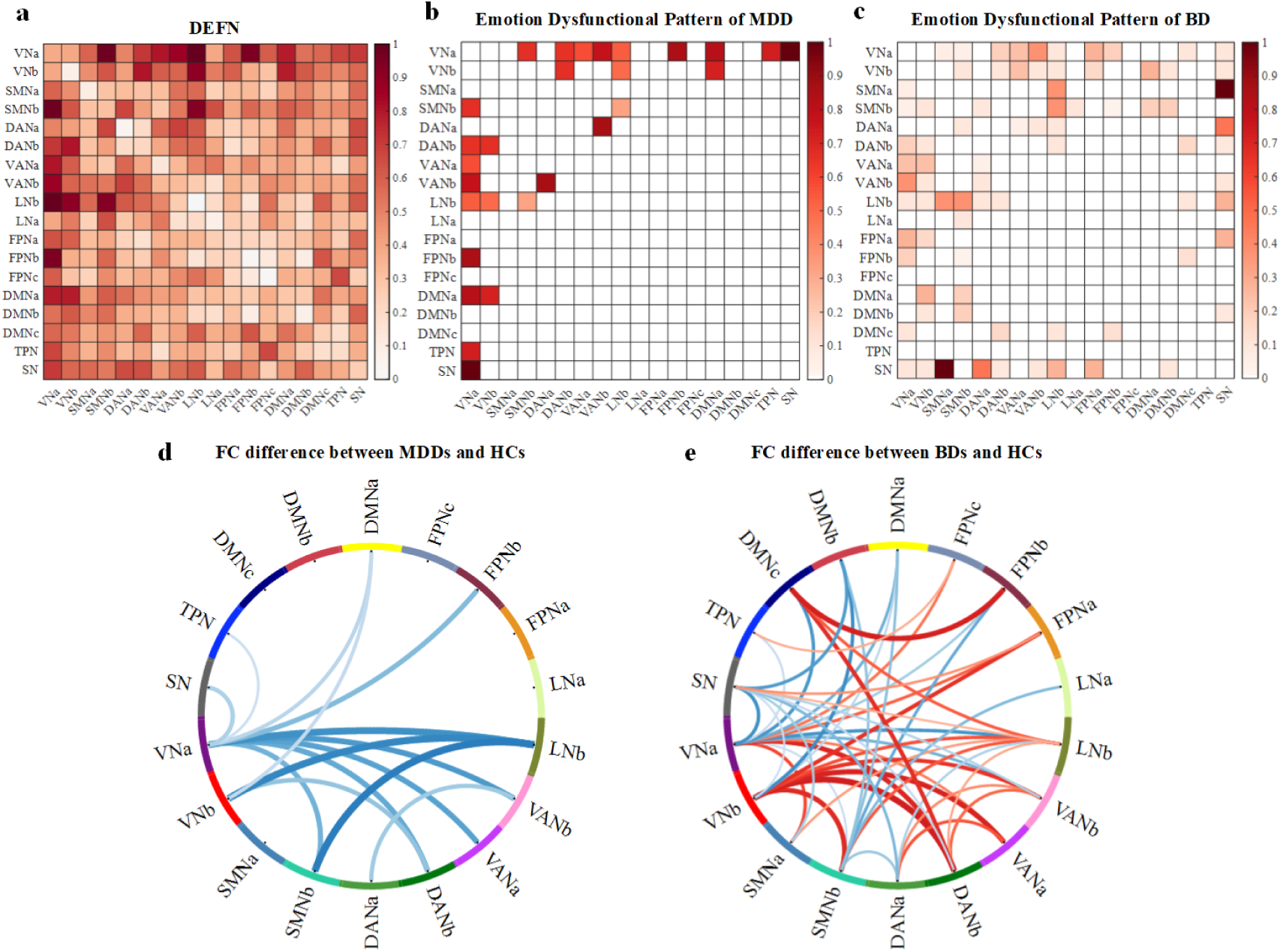
Identified DEFN and emotion dysfunctional patterns of MDD and BD patients. a, Identified DEFN based on the optimal model. Each square indicates a network weight, with the color gradient representing the magnitude: the darker the color, the greater the network weight. b, DEFN used by the optimal model trained on the MDD dataset. c, DEFN used by the optimal model trained on the BD dataset. The colors within the squares represent networks selected as DEFN indices, while white indicates networks that were not selected. Each square indicates a network importance, with the color gradient representing the magnitude: the darker the color, the greater the network importance. d, In the MDD dataset, the difference between the FCs of the MDD group and the HCs is shown, with blue indicating that this network connection is weaker in the MDD group compared to the HC group. e, In the BD dataset, the difference between the FCs of the BD group and those of the HC group is shown. Blue indicates that the BD group has fewer connections in this network compared to the HC group, while red indicates that the BD group has more connections than the HC group. The thickness of the line represents the difference, with thicker lines indicating a larger difference and vice versa.

**Fig. 4:**
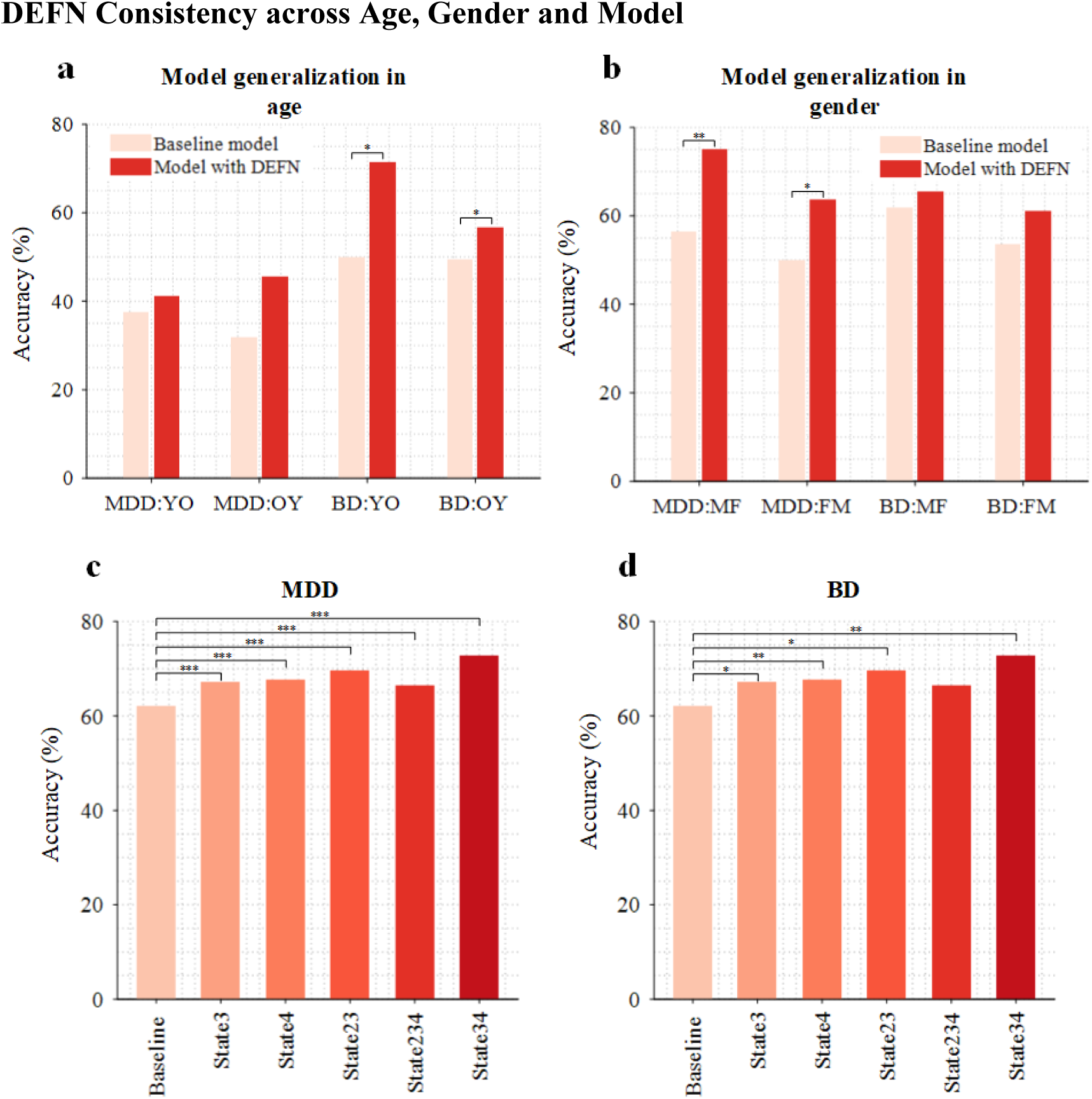
Results of DEFN consistency across age, gender, and model. a, Model generalization in age in MDD and BD datasets. YO indicates that the model is trained using data from the youth group and validated using data from the elderly group. OY, on the other hand, represents the opposite arrangement. b, Model generalization in gender in MDD and BD datasets. MF refers to a model trained on male-based data and validated on female data, while FM represents the opposite case. c, Classification accuracies of the models with DEFNs in the MDD dataset. d, Classification accuracies of the models with DEFNs in the BD dataset. Models based on state3, state4, state23, state234, and state34 were trained from the FC masked with DEFN information.

**Table 1.** Models’ performance for distinguishing happiness and sadness and significant differences (P value, corrected) of model’s performance between state34 and other states.

| Model | Accuracy | P value | Model | Accuracy | P value |
| --- | --- | --- | --- | --- | --- |
| State1 | 58.93±31.84% | 0.0001 | State24 | 80.20±22.53% | 0.1361 |
| State2 | 73.82±19.48% | 0.0027 | **State34** | **83.99±18.56%** | **-----** |
| State3 | 81.93±19.78% | 0.3750 | State123 | 73.21±24.22% | 0.0060 |
| State4 | 82.25±20.82% | 0.3750 | State124 | 76.03±20.75% | 0.0070 |
| State12 | 70.16±24.59% | 0.0021 | State134 | 71.67±25.91% | 0.0021 |
| State13 | 72.54±30.03% | 0.0190 | State234 | 84.25±19.06% | 0.8875 |
| State14 | 72.34±24.27% | 0.0190 | State1234 | 75.60±21.56% | 0.0085 |
| State23 | 81.93±18.20% | 0.4189 |  |  |  |

### Emotion Dysfunctional Patterns of MDD and BD

To investigate the neural patterns of emotional dysfunction in mood disorders, we explored two types of mental disorders: MDD and BD. Based on the prior information of the DEFN identified from state34 in healthy participants, this study established linear SVM models to separately distinguish MDDs and BDs from HCs with and without DEFN (referred to as the baseline model) in MDD and BD datasets. McNemar tests (i.e., paired chi-square tests) were employed to examine whether DEFN can significantly improve the model’s ability. The classification results and emotion dysfunctional patterns are presented in Fig.3.

The classification results in the MDD dataset indicate that the model incorporating the DEFN from the top 14 networks achieved the best performance, achieving a classification accuracy of 72.78%. Detailed classification results can be found in Supplementary TABLE S1. Compared to the baseline model (classification accuracy: 62.03%), the best-performing model exhibited a significant performance improvement of 10.75% (p<0.05, corrected). Additionally, the network pairs that contribute to distinguishing MDDs from HCs (i.e, emotion dysfunctional pattern of MDD) include VNa between-network pairs (DANb, VAN, LNb, FPNb, DMNa, TPN, SN), VNb between-network pairs (DANb, LNb, DMNa), SMNb-LNb, and DANa-VANb. These findings underscore the importance of these connections in the neural mechanisms of depression.

The classification results in the BD dataset show that the model incorporating the DEFN from the top 45 networks achieved the best performance, achieving a classification accuracy of 69.72%. Detailed classification results are provided in Supplementary TABLE S2. Compared to the baseline model (classification accuracy: 57.80%), the best-performing model exhibited a significant performance improvement of 9.92% (p<0.05, corrected). The network pairs that contribute to distinguishing BDs from HCs include not only those found in MDD but also VNb between-network pairs (SMNb, VANa, VANb, FPNa, DMNb, SN), SMNb between-network pairs (DANa, LNa, FPNb, DMNa, DMNb), DMNc between-network pairs (VNa, DANb, LNb, FPNb), and SN between-network pairs (VNb, SMNa, DANa, DANb, VANb, LNb, FPNa, DMNb). These findings underscore the shared and divergent neural connectivity disruptions in MDD and BD, providing critical insights into the underlying mechanisms of emotional dysfunction.

### DEFN Consistency across Age, Gender and Model

In the MDD and BD datasets, patients vary widely in age and include male and female genders. To verify the model’s generalization across different age groups, we divided the dataset into a young group and an older group. Individuals under the age of 35 were classified as the young group, while those over the age of 35 were classified as the middle-aged and older group. We first trained both the DEFN-based model and the baseline model on the young group and tested them on the old group. Then, we reversed the process by training the models on the old group and testing them on the young group. Additionally, McNemar tests were employed to determine whether there was any significant difference in performance between the baseline model and the model with DEFN. Additionally, on the emotional dataset, we obtained the optimal model from the feature combination of State34, as well as a second-best model with no statistically significant difference from the optimal model. To validate whether the DEFN constructed based on these second-best models could also effectively represent brain activity associated with mood disorders, we applied the DEFN derived from these models to the mood disorder dataset and evaluated its classification performance. Finally, the similarity between these DEFNs was assessed.

The results indicate that the model incorporating DEFN consistently exhibits significantly better generalization ability for age compared to the baseline model. In the MDD dataset, the classification accuracies of the model incorporating DEFN and the baseline model were 43.32% and 34.66%, respectively. Similarly, in the BD dataset, the classification accuracies were 64.11% and 49.69%, respectively. For gender generalization, we applied the same cross-group validation approach, training the models on one gender group and testing them on the other. The model incorporating DEFN also demonstrates superior generalization performance across gender. In the MDD dataset, it achieved a classification accuracy of 69.32%, outperforming the baseline model (53.26%). In the BD dataset, the classification accuracies for the DEFN-based model and the baseline model were 63.28% and 57.76%, respectively. These findings confirm that the DEFN-based model consistently outperforms the baseline model in generalization across both age and gender, reinforcing its robustness in distinguishing MDD and BD from healthy controls across diverse patient subgroups. All models with DEFN significantly outperformed the baseline model in the MDD dataset. In the BD dataset, all models, except the one constructed based on DEFN from State234, significantly outperformed the extreme model. The DEFNs constructed from the optimal model and second-best models exhibit strong correlations, with the correlation between any two DEFNs being statistically significant (Supplementary Fig S1).

## Discussion

To improve the development of novel diagnostic biomarkers for mood disorders and to investigate the emotional dysfunctional brain networks in mood disorders, this study developed and tested a novel two-stage approach using naturalistic movie paradigms to evoke ecological-valid and prolonged experiences of happiness and sadness in healthy participants and subsequently determining the applicability of this signature as neuromarker for mood disorders. By decoding the dynamic network-level signatures of these emotional states, we identified the Divergent Emotional Functional Network (DEFN), which characterizes the neural signature underlying positive and negative emotional experiences relevant to the core symptoms of mood disorders. Leveraging prior information from DEFN, we explored the emotional dysregulation patterns in two common mental disorders: MDD and BD. Specifically, based on the sliding window technique and clustering methods, four brain functional states reflecting dynamic emotional signatures were measured. The upper triangular elements of states were extracted as features and linear SVM models were built to identify DEFN. The optimal model, achieving a classification accuracy of 83.99%, demonstrated a precise ability to discriminate between two core emotional states: happiness and sadness. Furthermore, FC features masked with DEFN information, as well as whole-brain FC features, were separately extracted from MDD and BD datasets for constructing models. The inclusion of spatiotemporal information from the DEFN signature led to notable improvements of 10.75% and 9.92% in classification performance using resting-state data, effectively distinguishing MDD and BD patients from HCs. This enhancement underscores the importance of DEFN in identifying emotion dysregulation patterns in individuals with mood disorders.

### DEFN Reflecting Naturalistic Emotional Experiences

The decoding models trained to discriminate sad and happiness and sadness in healthy individuals indicate that high network weights in DEFN are primarily distributed among network pairs related to VNa, VNb, LNb, and SN. Previous studies have shown that emotions are associated with multiple regions of the cerebral cortex and subcortex with distributed patterns of activity and connectivity underlying complex emotional processes (Banks, *et al*., 2007, Čeko, *et al*., 2022, Johnstone, *et al*., 2007, Liu, *et al*., 2024). In addition to the reported involvement of individual brain regions, the generation and regulation of emotions involve complex networks composed of different brain areas and their interaction (Saarimäki, *et al*., 2022, Xu, *et al*., 2023, Zhou, *et al*., 2023). For instance, studies have consistently reported the involvement of the DMN in the neural representation of emotions (Buckner, *et al*., 2008). The DMN typically includes the medial prefrontal cortex (mPFC), posterior cingulate cortex (PCC), inferior parietal lobule (IPL), hippocampus, and other memory-related regions (Buckner, *et al*., 2008). Xu et al. found that using FC between multiple brain regions and networks enables more accurate decoding of happiness and sadness emotional experiences compared to using FC within a single network (Xu, *et al*., 2023). These findings indicate that emotions are encoded within the complex patterns of brain activity and interactions.

The complexity of emotional dysregulations makes the precise diagnosis and early intervention of mood disorders challenging and difficult. One possible reason is the existence of distinct neural representations for different emotions, with various subtypes of emotional disorders exhibiting dysregulation of specific emotions. A key symptom of depression represents anhedonia, probably related to dysfunctional reward processing (Bore et al., 2024; Bore et al., 2024) and alterations in the ventral striatum and associated areas. These regions form a network critically involved in pleasurable experiences, including the nucleus accumbens (Mahler, *et al*., 2007), the ventral pallidum (Ho and Berridge, 2013), and limbic areas of the prefrontal cortex (PFC), particularly the orbitofrontal cortex (OFC), anterior cingulate cortex (ACC), insular cortex (Berridge and Kringelbach, 2011) (Berridge and Kringelbach, 2013), and DMN. Our study also found that these brain regions are involved in the processing of happiness, and these regions are included in the LNb and DMNa. In addition, negative emotions such as sadness have been consistently reported to be primarily processed in subcortical regions, with additional contributions from visual areas (Pourtois and Vuilleumier, 2006) (Jaworska, *et al*., 2015) (Ramirez-Mahaluf, *et al*., 2018). The decoding results of this study also revealed that the connections between the visual regions, subcortical areas, and other networks are involved in the processing and integration of negative emotions. Considering that different mood disorders primarily involve changes in happiness and sadness, this study explores the impact of the differences between emotion of happiness and sadness on the identification of various mood disorders.

### Enhanced decoding of mood disorders using the DEFN as compared to the baseline model

This study demonstrates that incorporating the DEFN model with resting-state data leads to improved classification accuracy compared to the baseline model. Specifically, the DEFN model achieved a classification accuracy of 72.78% for identifying MDD patients, compared to 62.03% with the baseline model. Similarly, the DEFN model classified BD patients with an accuracy of 69.72%, outperforming the baseline model’s 57.80%. Compared to other classification results based on resting-state functional connectivity, the classification accuracy of our baseline model is similar. For instance, a study by Gallo et al. used rs-fMRI data from the REST-meta-MDD (N=2338) and PsyMRI (N=1039) consortia, along with SVM, to classify FC features of MDDs and HCs (Gallo, *et al*., 2023). The results showed that the model achieved an average classification accuracy of 61% in identifying MDDs from HCs, which is similar to the performance of the baseline model in our study. Another study by Dai et al. using a Transformer-Encoder model achieved an average classification accuracy of 68.61% on a dataset consisting of 832 MDDs and 779 HCs (Dai, *et al*., 2024). The lower classification accuracy of our baseline model may be due to the fact that we employed a more stringent leave-one-subject-out cross-validation method, whereas Dai’s study used a more relaxed five-fold cross-validation approach. Nevertheless, our DEFN model still achieved a higher classification accuracy than this model, exceeding it by 4.17%. Compared to other studies, our DEFN model improved accuracy by 4.17% to 11.78%, which holds significant research value for the clinical diagnosis of MDD patients (Dai, *et al*., 2024, Gallo, *et al*., 2023).

### Emotion Dysfunctional Pattern of MDD

This study accurately identifies patients with MDD from healthy individuals based on the functional activity of the visual network-attention network, prefrontal-visual network, and limbic network-visual network. And the connections between these networks in MDDs were generally weaker than in HCs, indicating reduced information interaction between different brain regions in MDDs. Previous studies have demonstrated patients with major depressive disorder and individuals with higher levels of depression exhibit functional impairments in multiple emotion-processing domains during resting-state, including the connection between the VN, DMN, VAN and PFC (Xu, *et al*., 2020) (Liu, *et al*., 2023) (Xu, *et al*., 2022). Studies have shown that patients with MDD exhibit attention deficits (Canario, *et al*., 2021) (Dai, *et al*., 2023). Through EEG microstate analysis, Wen et al. revealed that the probability of transition from microstate C to microstate D reflects anhedonia and attention deficits in the MDD group. Moreover, the temporal variability of the DAN in MDD patients was significantly lower compared to healthy controls (Wen, et al., 2023). Our findings indicate reduced connectivity between the visual and attention networks, which may suggest that deficits in the attention network of MDD patients lead to diminished visual focus, reflecting a decreased interest in exploring external stimuli.

Several of the connections identified encompassed connections between the lateral PFC (lPFC), mPFC, and VNa, suggesting that emotional dysfunctions in MDD may related to the well describe function of these regions in emotional processing and dysfunction. For instance, a meta-analysis reported that MDD might be associated with abnormal FC in the PFC, ACC, temporal lobe, and basal ganglia (Li, *et al*., 2022). The PFC is widely reported to be closely related to emotions and emotion regulation, including the lPFC (VANb and FPNb) and mPFC (DMNa). Previous task-based fMRI activation studies have reported insufficient activation of the lPFC in treatment-naïve MDD patients during social cognitive processes (Chen, *et al*., 2023) (Liu, *et al*., 2021), which normalized after antidepressant treatment (Siegle, *et al*., 2007). In a study using rsfMRI to classify HCs and MDDs, it was found that the ACC and lPFC were the most discriminative features during the classification process (Bondi, *et al*., 2023). This indicates that the lPFC encodes emotional processing and information integration, and that functional dysregulation in these regions contributes to MDDs. The mPFC, as a key region of the DMN, is associated with internal representations of self and reward value (Moran, *et al*., 2006). Furthermore, FC within the DMN is negatively correlated with the degree of self-rumination, suggesting that lower FC in the DMN is associated with higher levels of rumination, which is linked to a negative bias in emotional stimuli and impaired emotional self-regulation (Peng, *et al*., 2021).

### Emotion Dysfunctional Pattern of BD

Emotion dysfunctional patterns in BD include not only the connections typically associated with MDD but also involve additional brain networks (mainly including the connections between the VN, VAN, DAN, LNb, and SN), suggesting that BD may encompass broader and more complex abnormal neural activity. The LNb includes the lateral and medial OFC, which are major sites for the integration of multiple sensory modalities (Cole, *et al*., 2012) (Ochsner and Gross, 2005). We found abnormal FC between the OFC and the visual cortex and sensorimotor cortex in both MDD and BD. This suggests that the OFC integrates visual information and regulates top-down sensory-motor areas. Additionally, the OFC is a key brain region for human emotions, and its damage can impair subjective emotional states, emotional responses to stimuli such as facial and vocal expressions, and social behavior (Rolls, 1996) (Hornak, *et al*., 2003). This implies that the reduced FC in the OFC found in this study represents impaired emotional states, leading to behaviors such as slow speech and body movements in MDD patients, while the increased FC in the OFC might represent the heightened euphoria and energy experienced by BD patients during manic episodes. The study by Lu et al. utilized leave-one-subject-out cross-validation to differentiate HCs from BDs. The results demonstrated that FC between the right OFC and the left putamen identified BDs with an accuracy of 84.91%. (Lu, *et al*., 2021). Although the research methods employed by Lu et al. have certain data leakage limitations, the connectivity features between the OFC and the putamen were derived through traditional statistical analyses, which hold a certain degree of reference value.

Consistent with previous research, we found decreased connectivity between the SN and the visual cortex, sensory cortex, and attention networks. This may indicate that BD patients exhibit impaired attention and integration of visual and sensory-motor information during euphoric states, leading to abnormal perception and response to external stimuli (Phillips, *et al*., 2008) (Houenou, *et al*., 2011) (Öngür, *et al*., 2010). Research has shown that in BD patients, activation of the SN and PFC increases in response to extreme fear and mild pleasure (Lawrence, *et al*., 2004). This is consistent with our study, where enhanced connectivity between the SN and prefrontal cortex may suggest that these patients experience heightened sensitivity to positive and negative emotional stimuli, potentially contributing to increased emotional instability. A study utilizing rsFC (resting-state functional connectivity) features to classify MDD and BD highlighted the significance of subcortical regions, particularly the caudate nucleus, in distinguishing between the two disorders. Based on the rsFC of subcortical regions, MDD and BD patients were successfully classified with an accuracy of 88.00% (Jie, *et al*., 2018). This suggests that subcortical regions are critical functional brain features for differentiating MDD from BD patients.

## Methods

### Datasets

#### Self-acquired Healthy Dataset

A total of 52 healthy right-handed subjects (male/female: 26/26; age: 19 to 32 years old, 23.69±2.36; with normal or corrected-to-normal vision) from Shenzhen University were recruited by an advertisement for this experiment. Exclusion criteria for subject recruitment included neurological or psychiatric diagnosis, heavy alcohol consumption within the past six months, cardiovascular disease, severe visual impairment, and placement of metals in the body. All the subjects signed the informed consent before starting the experiment, and the experiment was approved by the Ethics Committee of the Health Science Center, Shenzhen University. The experiment was in line with the latest version of the Declaration of Helsinki. After quality control, data from one subject (female; 19 years old) was excluded due to excessive head motion (the exclusion criteria outlined in the data preprocessing section below). Details of the stimuli and experimental paradigm, as well as data acquisition procedures, are provided in the Supplementary Materials under the “Self-acquired Healthy Dataset” section.

#### Public MDD Dataset

The MDD dataset was collected from the multi-site, multi-disorder resting-state magnetic resonance image database created by Tanaka’s team (Tanaka, *et al*., 2021). Tanaka’s team has published 4 datasets: 1) the SRPBS Multi-disorder Connectivity Dataset 2), the SRPBS Multi-disorder MRI Dataset (restricted), 3) the SRPBS Multi-disorder MRI Dataset (unrestricted), and 4) the SRPBS Traveling Subject MRI Dataset. We employed the SRPBS Multi-disorder MRI Dataset (unrestricted) for the subsequent analysis. The SRPBS Multi-disorder MRI Dataset (unrestricted) consists of resting-state functional and structural MRI images of patients with 8 types of mental disorders (MDD, autistic spectrum disorders, obsessive-compulsive disorder, schizophrenia, pain, stroke, bipolar disorder, dysthymia, and others) as well as healthy participants. The dataset contains a total of 791 healthy participants and 619 patients from 11 sites. To eliminate the influence of different sites, we chose data from the same site with the highest number of healthy participants and MDD patients for analysis. The site with the highest number of participants had 174 individuals, including 111 healthy participants and 63 MDD patients. After preprocessing, data containing excessive head movements were excluded, resulting in 158 participants (58 MDDs and 100 HCs) for subsequent analysis. Details of data acquisition procedures are provided in the Supplementary Materials under the “Public MDD Dataset” section.

#### Self-acquired BD Dataset

The participants in this study comprised 61 euthymic and medicated BD patients (30 males, 31 females, age: 31.75±8.12 years) and 51 HCs (25 males, 26 females, age: 30.06±7.23 years). All BD patients were recruited from outpatient and inpatient departments at Shenzhen Mental Health Centre, and HCs were recruited by advertisement. The procedures of this experiment were approved by the Human Research Ethics Committee of Shenzhen Mental Health Center, and all participants in this experiment signed an informed consent form. After preprocessing, data containing excessive head movements were excluded, resulting in 109 participants (59 BDs and 50 HCs) for subsequent analysis. Details of data acquisition procedures are provided in the Supplementary Materials under the “Self-acquired BD Dataset” section.

### fMRI Preprocessing

All data were subjected to the following pre-processing steps. The functional images were preprocessed using SPM (Friston, 2003) and DPARSF (Yan, *et al*., 2016) in MATLAB. Given that the scanner included dummy scans to stabilize the magnetic field, the first five volumes of each time series were not discarded. However, BD data didn’t include dummy scans, thus the first five volumes of each time series were discarded. The structural images were first stripped of the skull and segmented into gray matter (GM), white matter (WM), and cerebrospinal fluid (CSF) based on the results of the stripped skull. Then, all the functional images of each episode were aligned with the first volume of functional images using six head-motion parametric linear transformations, and the functional images of each subject were coregistered with the structural images. To remove the linear drift and reduce the interference of head movements and other physiological signals, nuisance covariate regression was conducted using the Friston 24-parameter model. The functional images were then normalized to the Montreal Neurological Institute (MNI) space. A Gaussian kernel of 6 mm (FWHM) was adopted for spatial smoothing and bandpass filters of 0.008-0.15 in the healthy dataset and 0.01-0.1 Hz in MDD and BD datasets were conducted for eliminating the low-frequency drift and high-frequency noise and improving the signal-to-noise ratio of the BOLD signal (Wang, *et al*., 2017) (Meer, *et al*., 2020).

Data with excessive head motion were discarded according to the following exclusion criteria: (1) In the healthy dataset, an episode with a maximum translation of more than 2 mm or rotation of more than 2°; In MDD and BD datasets, a run with a maximum translation of more than 3 mm or rotation of more than 3°; (2) for one subject, if more than 2 episodes (total 6 episodes) were discarded, data from this subject would be fully excluded.

### Brain Parcellation

We applied the Schaefer parcellation (Schaefer, *et al*., 2018) covering the whole brain cortex with 400 Regions of Interest (ROIs), and the Tian atlas (Tian, *et al*., 2020) covering the brain subcortex with 32 ROIs, to extract the BOLD time series. For each of 432 ROIs, we averaged the BOLD time series across all voxels within each ROI. Schaefer parcellation integrated both local gradient and global similarity from rest-state and task-state FC. According to the clearly defined coordinates of the location of structural subdivisions in the whole-brain cortex, the parceled ROIs could be categorized into 17 networks including Visual Network (VN: VNa, VNb), SomatoMotor Network (SMN: SMNa, SMNb), Dorsal Attention Network (DAN: DANa, DANb), Ventral Attention Network (VAN: VANa, VANb), Limbic Network (LN: LNa, LNb), FrontoParietal Network (FPN: FPNa, FPNb, FPNc), Default Mode Network (DMN: DMNa, DMNb, DMNc), and Temporal Parietal Network (TPN).

Besides, consistent with previous studies (Luppi and Stamatakis, 2021) (Luppi, *et al*., 2022), 32 subcortical ROIs were also considered based on the recently developed Melbourne subcortical functional parcellation atlas, which was obtained based on resting-state and task-state functionally connectivity and was consistent with the parcellation methodology in Schaefer 400. The 32 subcortical ROIs covered 7 subcortical regions, including the hippocampus, thalamus, amygdala, caudate nucleus, putamen, and globus pallidus. All the 32 ROIs were grouped into the Subcortex Network (SN). In total, 18 networks (17 cortical networks and 1 subcortical network) were included reflecting a broad and more fine-grained level of organization, respectively.

### States Estimation on Healthy Participants

For the extracted BOLD time series, the sliding window method (time window length:30 TRs, sliding step: 2 TRs) was used to estimate the FC matrices over time. Specifically, the entire BOLD time series was split into several time series segments by a fixed time window (30 TRs) length and sliding step (2 TRs). The Pearson correlation coefficient was subsequently utilized to estimate the correlation between each pair of 432 ROIs across each time window, generating a matrix (286×432×432) describing the dFC for each movie segment. Here, 286 represents the time length of dFC, and 432 represents the number of ROIs. Additionally, Fisher’s Z-transform was applied to improve the normality of the correlation coefficients.

The k-means clustering method was employed to identify recurrent brain states under happiness and sadness based on dFC. Before conducting the clustering analysis, we determined the optimal number of clusters using the elbow method as it significantly influences the clustering results. In the clustering analysis, the L1 distance was employed to gauge the similarity between two dFC matrices across all participants by considering their upper triangular elements. This distance metric is recognized as an effective measure for high-dimensional data. Specifically, the K-means clustering was performed in two steps. In the first step, a subset of dFCs was selected as input. This subset comprises the windows corresponding to the local maxima of FC variance. The L1 distance metric was employed to randomly determine the initial clustering center-of-mass position, which was then repeated 100 times. The resulting outcome preserves the center-of-mass position of the clusters. In the second step, the k-means clustering was carried out on the episodes of all subjects. The centroids of the first step clustering were set as the initial points, and the number of iterations for k-means was set to 1000. Finally, all dFC matrices were clustered into 4 states according to the elbow method.

### DEFN Identification on Healthy Participants

Based on the states, the connectivity strength within each state and combinations of connectivity strengths across different states were extracted as features to decode happiness and sadness experiences, thereby identifying DEFN from the perspective of neural activity. Here, we constructed 15 decoding models, including models built based on state1, state2, state3, state4, state12, state13, state14, state23, state24, state34, state123, state124, state134, state234, and state1234. Specifically, for individual states (taking state1 as an example) and each episode, the averaged matrix of state1 was computed by averaging all state1 matrices. Subsequently, the upper triangular elements of the state1 matrix were extracted as features for decoding happiness and sadness. For feature combinations across states (taking state12 as an example), the upper triangular elements of the averaged state1 matrix and state2 matrix were extracted, and these connectivity features were concatenated as features. Leave-one-subject-out cross-validation and relief feature selection methods were employed to train linear SVM models.

For the constructed decoding models, classification accuracy was employed to estimate the decoding performance of the models, and the model with the highest classification accuracy was regarded as the optimal model. T-tests and false discovery rate (FDR) correction were applied to determine whether there was a significant difference in classification accuracy between the optimal model and other models. Based on the optimal model, features consistently emerged across per-fold cross-validation and were extracted as stable features, obtaining the distribution of the stable features in each state. Subsequently, the union of these stable features across states was extracted to evaluate the network weights (referred to as DEFN) by calculating the ratio of the number of stable features to the total number of features within each network pair.

### Exploration of Emotion Dysfunctional Patterns of MDD and BD

To investigate the neural patterns of emotional dysfunction in mood disorders, we explored two types of mental disorders (MDD and BD) based on the prior information of the DEFN identified from healthy participants. Firstly, the network weights were sorted in descending order, and the top n networks were sequentially chosen as masks (for example, selecting the top-ranked network as the mask in the first turn, and the top two networks in the second turn), resulting in a series of DEFN masks. Subsequently, these DEFN masks were individually applied to the MDD dataset and the BD dataset to extract important features from whole-brain FC, building classification models that identify MDDs or BDs from HCs. Furthermore, the whole-brain FC features from MDD or BD datasets were also extracted to construct the baseline model, aiming to validate whether the performance of models with DEFN masks was improved. Then, McNemar tests (i.e., a paired chi-square test) were employed to determine whether there was any significant difference in performance between the baseline model and the model with DEFN mask. In addition, we calculated the importance of each network pair of DEFN-based model for classifying mood disorder through a permutation test. The specific steps are as follows: (1) For each network pair’s features, randomly shuffle these features across all subjects; (2) Using the shuffled features and leave-one-out cross-validation, train the model and evaluate its performance; (3) Repeat steps (1) and (2) 50 times to obtain the average classification accuracy across 50 iterations; (4) Use the classification accuracy of the model built with the original features and subtract the classification accuracy of the model built with the shuffled features as the importance measure for the network pair; (5) Repeat the above steps for all network pairs.

### Validation of DEFN Consistency across Age, Gender and Model

In the MDD and BD datasets, patients vary widely in age and include male and female genders. To verify the model’s generalization across different age groups, we divided the dataset into a young group and an older group. Individuals under the age of 35 were classified as the young group, while those over the age of 35 were classified as the middle-aged and older group. The reason for choosing 35 years as the age cutoff is that all subjects in the emotional dataset are younger than 35 years. Additionally, considering the large age difference between the subjects in the MDD and BD datasets, the BD dataset is predominantly composed of adolescents, while the MDD dataset has a majority of elderly participants. To balance the number of participants in the two groups from both datasets, 35 years was selected as the age cutoff. We first trained both the DEFN-based model and the baseline model on the young group and tested them on the old group. Then, we reversed the process by training the models on the old group and testing them on the young group. Additionally, McNemar tests were employed to determine whether there was any significant difference in performance between the baseline model and the models with DEFNs. Additionally, on the emotional dataset, we obtained the optimal model from the feature combination of State34, as well as a second-best model with no statistically significant difference from the optimal model. To validate whether the DEFN constructed based on these second-best models could also effectively represent brain activity associated with mood disorders, we applied the DEFN derived from these models to the mood disorder dataset and evaluated its classification performance. Finally, the similarity between these DEFNs was assessed.

## Acknowledgments

This work was supported in part by the National Natural Science Foundation of China under Grant 62276169, in part by the Medical-Engineering Interdisciplinary Research Foundation of Shenzhen University under Grant 2024YG008, in part by the Shenzhen University-Lingnan University Joint Research Programme, in part by the Shenzhen-Hong Kong Institute of Brain Science-Shenzhen Fundamental Research Institutions under Grant 2023SHIBS0003, in part by the STI 2030-Major Projects 2021ZD0200500, and in part by the Open Research Fund of the State Key Laboratory of Brain-Machine Intelligence, Zhejiang University (Grant No. BMI2400008).

## Supplementary Materials

### Performance of Models with DEFN in MDD and BD datasets

#### Consistency of DEFNs constructed from optimal models

To validate the consistency of these DEFNs across optimal models derived from state34, state3, state4, state23, and state234, the Pearson correlation coefficient was applied to estimate the similarity across them. The results indicate that the DEFNs constructed from the optimal models exhibit strong correlations, with the correlation between any two DEFNs being statistically significant. What’s more, all models with DEFN significantly outperformed the baseline model in the MDD dataset. In the BD dataset, all models, except the one constructed based on DEFN from dFC234, significantly outperformed the extreme model.

**Fig S1:**
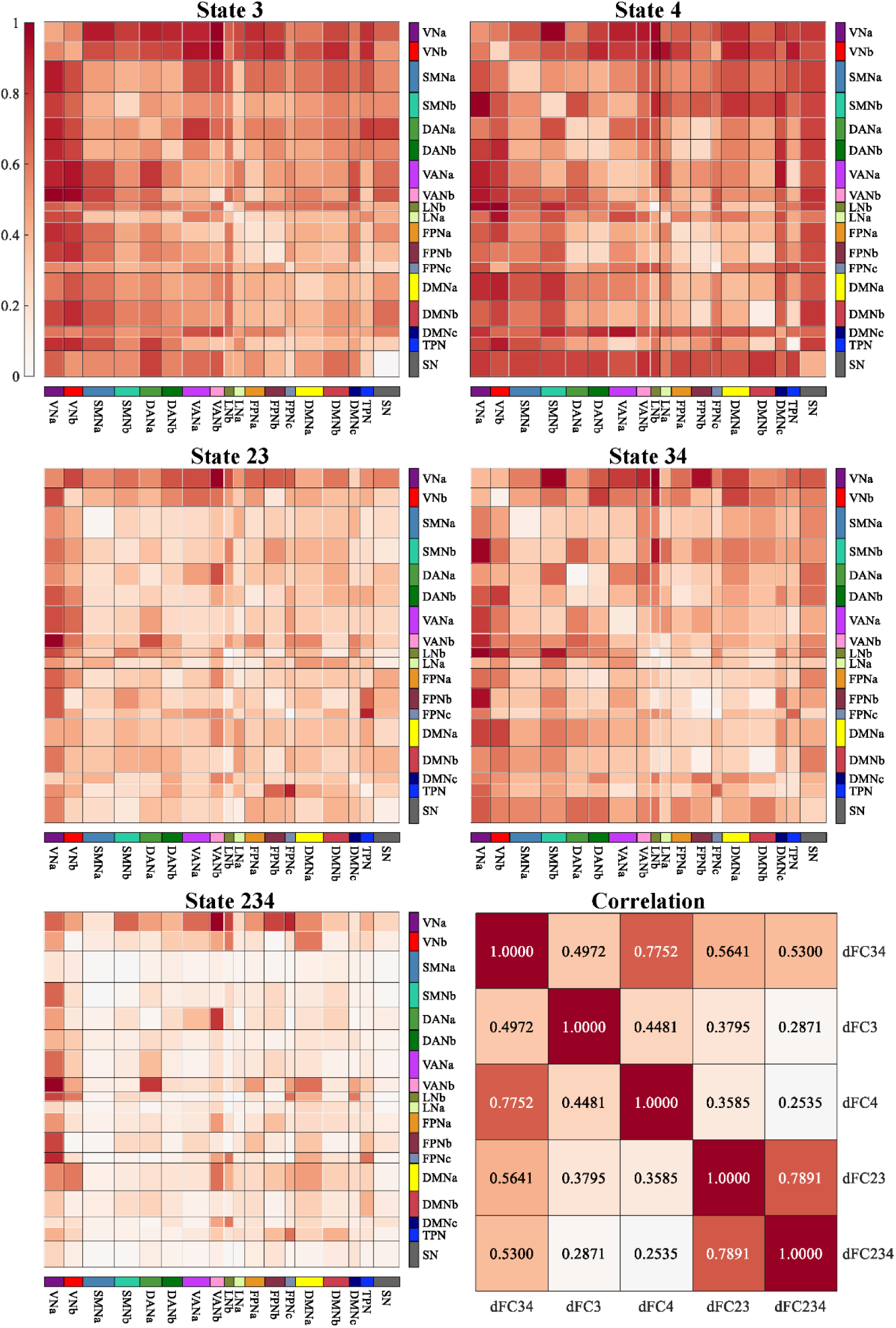
The identified DEFNs from State3, State4, State23, State234. Each network indicates the network weight, with the color gradient representing the magnitude: the darker the color, the greater the network weight. The Pearson correlation coefficient of the DEFNs obtained across state34, state3, state4, state23, and state234.

**TABLE S1:** Models’ performance built on State34 in MDD dataset.

| Model | Accuracy | Model | Accuracy | Model | Accuracy | Model | Accuracy | Model | Accuracy |
| --- | --- | --- | --- | --- | --- | --- | --- | --- | --- |
| Baseline | 62.03 | Top 34 | 63.29 | Top 68 | 63.29 | Top 102 | 60.76 | Top 136 | 61.39 |
| Top 1 | 57.59 | Top 35 | 62.03 | Top 69 | 63.29 | Top103 | 60.76 | Top 137 | 61.39 |
| Top 2 | 58.86 | Top 36 | 62.66 | Top 70 | 61.39 | Top 104 | 63.29 | Top 138 | 58.23 |
| Top 3 | 61.39 | Top 37 | 60.76 | Top 71 | 56.33 | Top 105 | 64.56 | Top 139 | 60.13 |
| Top 4 | 56.33 | Top 38 | 63.29 | Top 72 | 59.49 | Top 106 | 62.66 | Top 140 | 61.39 |
| Top 5 | 63.92 | Top 39 | 63.29 | Top 73 | 61.39 | Top 107 | 62.66 | Top 141 | 61.39 |
| Top 6 | 65.82 | Top 40 | 58.86 | Top 74 | 60.13 | Top 108 | 62.03 | Top 142 | 62.66 |
| Top 7 | 65.82 | Top 41 | 61.39 | Top 75 | 56.96 | Top 109 | 63.29 | Top 143 | 62.03 |
| Top 8 | 67.09 | Top 42 | 59.49 | Top 76 | 60.76 | Top 110 | 61.39 | Top 144 | 62.66 |
| Top 9 | 61.39 | Top 43 | 60.13 | Top 77 | 58.86 | Top 111 | 62.03 | Top 145 | 62.03 |
| Top 10 | 68.35 | Top 44 | 58.86 | Top 78 | 58.23 | Top 112 | 62.03 | Top 146 | 61.39 |
| Top 11 | 60.76 | Top 45 | 61.39 | Top 79 | 57.59 | Top 113 | 60.13 | Top 147 | 61.39 |
| Top 12 | 67.09 | Top 46 | 61.39 | Top 80 | 58.86 | Top 114 | 61.39 | Top 148 | 59.49 |
| Top 13 | 64.56 | Top 47 | 58.23 | Top 81 | 59.49 | Top 115 | 61.39 | Top 149 | 60.13 |
| Top 14 | 72.78 | Top 48 | 56.33 | Top 82 | 61.39 | Top 116 | 63.92 | Top 150 | 59.49 |
| Top 15 | 59.49 | Top 49 | 62.03 | Top 83 | 58.86 | Top 117 | 62.66 | Top 151 | 60.13 |
| Top 16 | 57.59 | Top 50 | 67.09 | Top 84 | 59.49 | Top 118 | 64.56 | Top 152 | 59.49 |
| Top 17 | 60.13 | Top 51 | 62.66 | Top 85 | 60.13 | Top 119 | 60.13 | Top 153 | 60.13 |
| Top 18 | 59.49 | Top 52 | 59.49 | Top 86 | 60.76 | Top 120 | 63.29 | Top 154 | 59.49 |
| Top 19 | 62.03 | Top 53 | 63.92 | Top 87 | 60.13 | Top 121 | 62.66 | Top 155 | 62.03 |
| Top 20 | 63.29 | Top 54 | 63.29 | Top 88 | 58.23 | Top 122 | 60.76 | Top 157 | 60.76 |
| Top 21 | 62.03 | Top 55 | 61.39 | Top 89 | 62.03 | Top 123 | 63.92 | Top 157 | 60.76 |
| Top 22 | 63.92 | Top 56 | 61.39 | Top 90 | 62.66 | Top 124 | 65.82 | Top 159 | 62.03 |
| Top 23 | 63.92 | Top 57 | 62.03 | Top 91 | 59.49 | Top 125 | 65.19 | Top 160 | 61.39 |
| Top 24 | 60.13 | Top 58 | 61.39 | Top 92 | 58.86 | Top 126 | 65.82 | Top 161 | 61.39 |
| Top 25 | 59.49 | Top 59 | 60.76 | Top 93 | 60.13 | Top 127 | 64.56 | Top 162 | 60.76 |
| Top 26 | 59.49 | Top 60 | 62.66 | Top 94 | 60.13 | Top 128 | 63.29 | Top 163 | 59.49 |
| Top 27 | 60.76 | Top 61 | 62.03 | Top 95 | 58.23 | Top 129 | 63.29 | Top 164 | 59.49 |
| Top 28 | 65.19 | Top 62 | 61.39 | Top 96 | 58.23 | Top 130 | 63.29 | Top 165 | 58.23 |
| Top 29 | 64.56 | Top 63 | 59.49 | Top 97 | 59.49 | Top 131 | 63.29 | Top 166 | 57.59 |
| Top 30 | 61.39 | Top 64 | 56.96 | Top 98 | 67.09 | Top 132 | 63.92 | Top 167 | 60.13 |
| Top 31 | 61.39 | Top 65 | 56.96 | Top 99 | 59.49 | Top 133 | 63.92 | Top 168 | 60.76 |
| Top 32 | 62.03 | Top 66 | 60.13 | Top 100 | 67.72 | Top 134 | 57.59 | Top 169 | 58.86 |
| Top 33 | 60.13 | Top 67 | 63.92 | Top 101 | 62.03 | Top 135 | 62.66 | Top 170 | 61.39 |

**TABLE S2:** Models’ performance built on State34 in BD dataset.

| Model | Accuracy | Model | Accuracy | Model | Accuracy | Model | Accuracy | Model | Accuracy |
| --- | --- | --- | --- | --- | --- | --- | --- | --- | --- |
| Baseline | 57.80 | Top 34 | 64.22 | Top 68 | 61.47 | Top 102 | 61.47 | Top 136 | 62.39 |
| Top 1 | 46.79 | Top 35 | 63.30 | Top 69 | 62.39 | Top 103 | 64.22 | Top 137 | 61.47 |
| Top 2 | 53.21 | Top 36 | 65.14 | Top 70 | 62.39 | Top 104 | 61.47 | Top 138 | 58.72 |
| Top 3 | 55.96 | Top 37 | 66.97 | Top 71 | 62.39 | Top 105 | 58.72 | Top 139 | 62.39 |
| Top 4 | 59.63 | Top 38 | 67.89 | Top 72 | 65.14 | Top 106 | 57.80 | Top 140 | 60.55 |
| Top 5 | 59.63 | Top 39 | 64.22 | Top 73 | 64.22 | Top 107 | 59.63 | Top 141 | 61.47 |
| Top 6 | 61.47 | Top 40 | 66.97 | Top 74 | 66.06 | Top 108 | 56.88 | Top 142 | 61.47 |
| Top 7 | 62.39 | Top 41 | 66.06 | Top 75 | 66.06 | Top 109 | 55.96 | Top 143 | 62.39 |
| Top 8 | 64.22 | Top 42 | 66.06 | Top 76 | 63.30 | Top 110 | 55.96 | Top 144 | 62.39 |
| Top 9 | 64.22 | Top 43 | 63.30 | Top 77 | 65.14 | Top 111 | 55.96 | Top 145 | 61.47 |
| Top 10 | 58.72 | Top 44 | 65.14 | Top 78 | 65.14 | Top 112 | 56.88 | Top 146 | 62.39 |
| Top 11 | 64.22 | Top 45 | 69.72 | Top 79 | 65.14 | Top 113 | 58.72 | Top 147 | 61.47 |
| Top 12 | 63.30 | Top 46 | 67.89 | Top 80 | 65.14 | Top 114 | 59.63 | Top 148 | 61.47 |
| Top 13 | 66.06 | Top 47 | 64.22 | Top 81 | 65.14 | Top 115 | 59.63 | Top 149 | 61.47 |
| Top 14 | 61.47 | Top 48 | 67.89 | Top 82 | 66.06 | Top 116 | 60.55 | Top 150 | 63.30 |
| Top 15 | 58.72 | Top 49 | 66.97 | Top 83 | 63.30 | Top 117 | 61.47 | Top 151 | 61.47 |
| Top 16 | 60.55 | Top 50 | 62.39 | Top 84 | 63.30 | Top 118 | 60.55 | Top 152 | 59.63 |
| Top 17 | 60.55 | Top 51 | 67.89 | Top 85 | 64.22 | Top 119 | 61.47 | Top 153 | 61.47 |
| Top 18 | 62.39 | Top 52 | 67.89 | Top 86 | 65.14 | Top 120 | 60.55 | Top 154 | 59.63 |
| Top 19 | 62.39 | Top 53 | 67.89 | Top 87 | 66.06 | Top 121 | 59.63 | Top 155 | 60.55 |
| Top 20 | 65.14 | Top 54 | 65.14 | Top 88 | 64.22 | Top 122 | 59.63 | Top 157 | 62.39 |
| Top 21 | 61.47 | Top 55 | 66.06 | Top 89 | 62.39 | Top 123 | 61.47 | Top 157 | 62.39 |
| Top 22 | 59.63 | Top 56 | 62.39 | Top 90 | 61.47 | Top 124 | 59.63 | Top 159 | 61.47 |
| Top 23 | 59.63 | Top 57 | 64.22 | Top 91 | 60.55 | Top 125 | 60.55 | Top 160 | 64.22 |
| Top 24 | 62.39 | Top 58 | 60.55 | Top 92 | 60.55 | Top 126 | 59.63 | Top 161 | 61.47 |
| Top 25 | 66.97 | Top 59 | 63.30 | Top 93 | 64.22 | Top 127 | 59.63 | Top 162 | 62.39 |
| Top 26 | 66.06 | Top 60 | 63.30 | Top 94 | 64.22 | Top 128 | 57.80 | Top 163 | 56.88 |
| Top 27 | 65.14 | Top 61 | 64.22 | Top 95 | 65.14 | Top 129 | 55.96 | Top 164 | 61.47 |
| Top 28 | 66.06 | Top 62 | 65.14 | Top 96 | 65.14 | Top 130 | 57.80 | Top 165 | 64.22 |
| Top 29 | 63.30 | Top 63 | 66.97 | Top 97 | 64.22 | Top 131 | 57.80 | Top 166 | 64.22 |
| Top 30 | 63.30 | Top 64 | 64.22 | Top 98 | 60.55 | Top 132 | 55.96 | Top 167 | 63.30 |
| Top 31 | 68.81 | Top 65 | 61.47 | Top 99 | 65.14 | Top 133 | 58.72 | Top 168 | 62.39 |
| Top 32 | 64.22 | Top 66 | 63.30 | Top 100 | 61.47 | Top 134 | 60.55 | Top 169 | 60.55 |
| Top 33 | 63.30 | Top 67 | 61.47 | Top 101 | 61.47 | Top 135 | 61.47 | Top 170 | 61.47 |

**TABLE S3:**
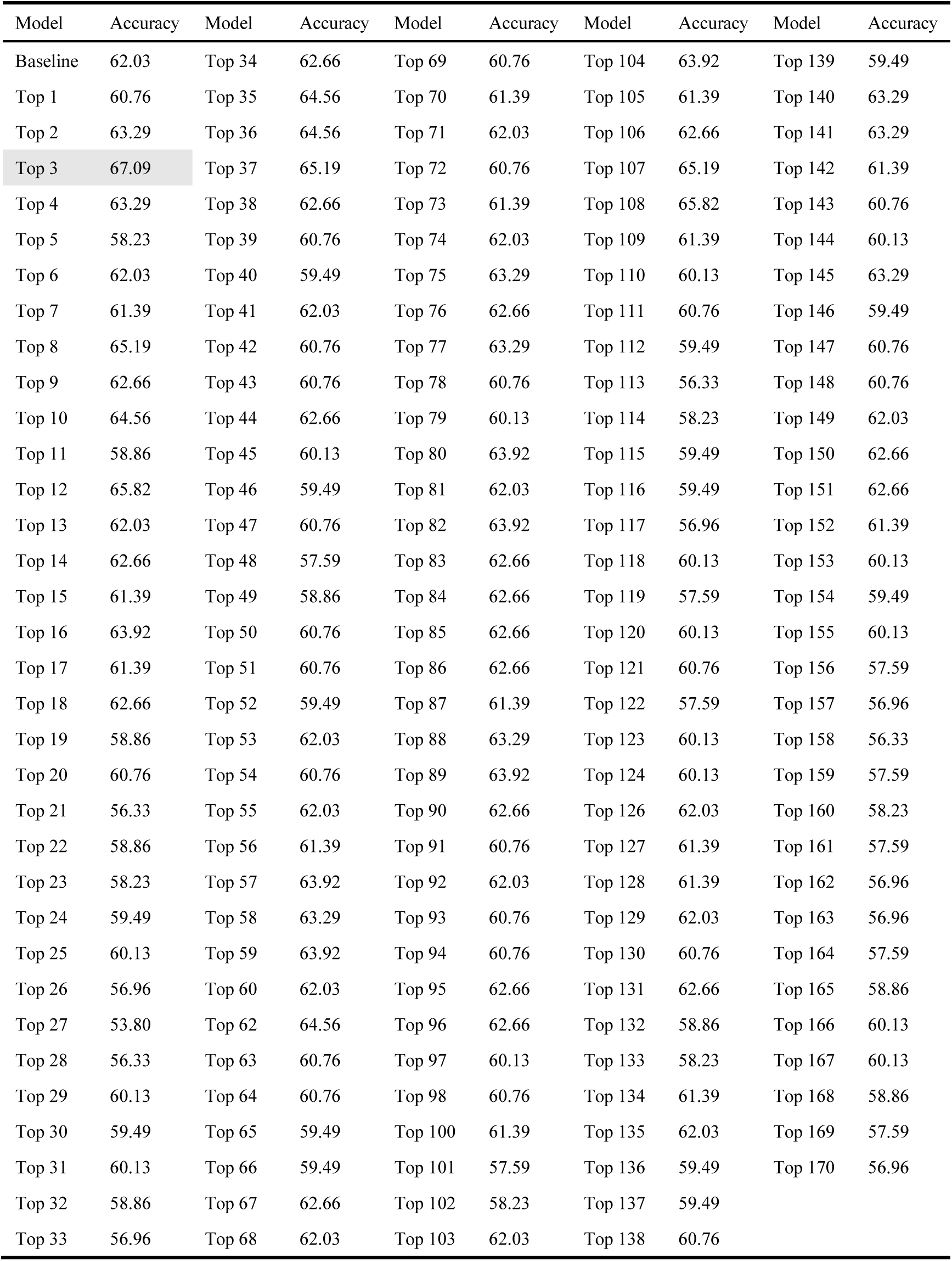
Models’ performance built on State3 in MDD dataset.

| Model | Accuracy | Model | Accuracy | Model | Accuracy | Model | Accuracy | Model | Accuracy |
| --- | --- | --- | --- | --- | --- | --- | --- | --- | --- |
| Baseline | 62.03 | Top 34 | 62.66 | Top 69 | 60.76 | Top 104 | 63.92 | Top 139 | 59.49 |
| Top 1 | 60.76 | Top 35 | 64.56 | Top 70 | 61.39 | Top 105 | 61.39 | Top 140 | 63.29 |
| Top 2 | 63.29 | Top 36 | 64.56 | Top 71 | 62.03 | Top 106 | 62.66 | Top 141 | 63.29 |
| Top 3 | 67.09 | Top 37 | 65.19 | Top 72 | 60.76 | Top 107 | 65.19 | Top 142 | 61.39 |
| Top 4 | 63.29 | Top 38 | 62.66 | Top 73 | 61.39 | Top 108 | 65.82 | Top 143 | 60.76 |
| Top 5 | 58.23 | Top 39 | 60.76 | Top 74 | 62.03 | Top 109 | 61.39 | Top 144 | 60.13 |
| Top 6 | 62.03 | Top 40 | 59.49 | Top 75 | 63.29 | Top 110 | 60.13 | Top 145 | 63.29 |
| Top 7 | 61.39 | Top 41 | 62.03 | Top 76 | 62.66 | Top 111 | 60.76 | Top 146 | 59.49 |
| Top 8 | 65.19 | Top 42 | 60.76 | Top 77 | 63.29 | Top 112 | 59.49 | Top 147 | 60.76 |
| Top 9 | 62.66 | Top 43 | 60.76 | Top 78 | 60.76 | Top 113 | 56.33 | Top 148 | 60.76 |
| Top 10 | 64.56 | Top 44 | 62.66 | Top 79 | 60.13 | Top 114 | 58.23 | Top 149 | 62.03 |
| Top 11 | 58.86 | Top 45 | 60.13 | Top 80 | 63.92 | Top 115 | 59.49 | Top 150 | 62.66 |
| Top 12 | 65.82 | Top 46 | 59.49 | Top 81 | 62.03 | Top 116 | 59.49 | Top 151 | 62.66 |
| Top 13 | 62.03 | Top 47 | 60.76 | Top 82 | 63.92 | Top 117 | 56.96 | Top 152 | 61.39 |
| Top 14 | 62.66 | Top 48 | 57.59 | Top 83 | 62.66 | Top 118 | 60.13 | Top 153 | 60.13 |
| Top 15 | 61.39 | Top 49 | 58.86 | Top 84 | 62.66 | Top 119 | 57.59 | Top 154 | 59.49 |
| Top 16 | 63.92 | Top 50 | 60.76 | Top 85 | 62.66 | Top 120 | 60.13 | Top 155 | 60.13 |
| Top 17 | 61.39 | Top 51 | 60.76 | Top 86 | 62.66 | Top 121 | 60.76 | Top 156 | 57.59 |
| Top 18 | 62.66 | Top 52 | 59.49 | Top 87 | 61.39 | Top 122 | 57.59 | Top 157 | 56.96 |
| Top 19 | 58.86 | Top 53 | 62.03 | Top 88 | 63.29 | Top 123 | 60.13 | Top 158 | 56.33 |
| Top 20 | 60.76 | Top 54 | 60.76 | Top 89 | 63.92 | Top 124 | 60.13 | Top 159 | 57.59 |
| Top 21 | 56.33 | Top 55 | 62.03 | Top 90 | 62.66 | Top 126 | 62.03 | Top 160 | 58.23 |
| Top 22 | 58.86 | Top 56 | 61.39 | Top 91 | 60.76 | Top 127 | 61.39 | Top 161 | 57.59 |
| Top 23 | 58.23 | Top 57 | 63.92 | Top 92 | 62.03 | Top 128 | 61.39 | Top 162 | 56.96 |
| Top 24 | 59.49 | Top 58 | 63.29 | Top 93 | 60.76 | Top 129 | 62.03 | Top 163 | 56.96 |
| Top 25 | 60.13 | Top 59 | 63.92 | Top 94 | 60.76 | Top 130 | 60.76 | Top 164 | 57.59 |
| Top 26 | 56.96 | Top 60 | 62.03 | Top 95 | 62.66 | Top 131 | 62.66 | Top 165 | 58.86 |
| Top 27 | 53.80 | Top 62 | 64.56 | Top 96 | 62.66 | Top 132 | 58.86 | Top 166 | 60.13 |
| Top 28 | 56.33 | Top 63 | 60.76 | Top 97 | 60.13 | Top 133 | 58.23 | Top 167 | 60.13 |
| Top 29 | 60.13 | Top 64 | 60.76 | Top 98 | 60.76 | Top 134 | 61.39 | Top 168 | 58.86 |
| Top 30 | 59.49 | Top 65 | 59.49 | Top 100 | 61.39 | Top 135 | 62.03 | Top 169 | 57.59 |
| Top 31 | 60.13 | Top 66 | 59.49 | Top 101 | 57.59 | Top 136 | 59.49 | Top 170 | 56.96 |
| Top 32 | 58.86 | Top 67 | 62.66 | Top 102 | 58.23 | Top 137 | 59.49 |  |  |
| Top 33 | 56.96 | Top 68 | 62.03 | Top 103 | 62.03 | Top 138 | 60.76 |  |  |

**TABLE S4:** Models’ performance built on State3 in BD dataset.

| Model | Accuracy | Model | Accuracy | Model | Accuracy | Model | Accuracy | Model | Accuracy |
| --- | --- | --- | --- | --- | --- | --- | --- | --- | --- |
| Baseline | 57.80 | Top 34 | 55.96 | Top 69 | 60.55 | Top 104 | 62.39 | Top 139 | 64.22 |
| Top 1 | 57.80 | Top 35 | 59.63 | Top 70 | 64.22 | Top 105 | 62.39 | Top 140 | 64.22 |
| Top 2 | 55.05 | Top 36 | 56.88 | Top 71 | 63.30 | Top 106 | 62.39 | Top 141 | 62.39 |
| Top 3 | 52.29 | Top 37 | 55.96 | Top 72 | 60.55 | Top 107 | 62.39 | Top 142 | 62.39 |
| Top 4 | 51.38 | Top 38 | 57.80 | Top 73 | 62.39 | Top 108 | 63.30 | Top 143 | 62.39 |
| Top 5 | 55.05 | Top 39 | 55.96 | Top 74 | 62.39 | Top 109 | 62.39 | Top 144 | 63.30 |
| Top 6 | 57.80 | Top 40 | 55.96 | Top 75 | 63.30 | Top 110 | 63.30 | Top 145 | 61.47 |
| Top 7 | 58.72 | Top 41 | 57.80 | Top 76 | 62.39 | Top 111 | 62.39 | Top 146 | 63.30 |
| Top 8 | 57.80 | Top 42 | 56.88 | Top 77 | 61.47 | Top 112 | 62.39 | Top 147 | 62.39 |
| Top 9 | 55.05 | Top 43 | 55.96 | Top 78 | 60.55 | Top 113 | 64.22 | Top 148 | 60.55 |
| Top 10 | 52.29 | Top 44 | 56.88 | Top 79 | 62.39 | Top 114 | 64.22 | Top 149 | 60.55 |
| Top 11 | 55.96 | Top 45 | 57.80 | Top 80 | 62.39 | Top 115 | 61.47 | Top 150 | 61.47 |
| Top 12 | 55.96 | Top 46 | 60.55 | Top 81 | 62.39 | Top 116 | 63.30 | Top 151 | 62.39 |
| Top 13 | 55.96 | Top 47 | 61.47 | Top 82 | 60.55 | Top 117 | 64.22 | Top 152 | 63.30 |
| Top 14 | 58.72 | Top 48 | 60.55 | Top 83 | 61.47 | Top 118 | 64.22 | Top 153 | 62.39 |
| Top 15 | 55.96 | Top 49 | 58.72 | Top 84 | 64.22 | Top 119 | 62.39 | Top 154 | 62.39 |
| Top 16 | 54.13 | Top 50 | 57.80 | Top 85 | 65.14 | Top 120 | 60.55 | Top 155 | 62.39 |
| Top 17 | 58.72 | Top 51 | 59.63 | Top 86 | 66.06 | Top 121 | 59.63 | Top 156 | 62.39 |
| Top 18 | 55.96 | Top 52 | 59.63 | Top 87 | 64.22 | Top 122 | 57.80 | Top 157 | 65.14 |
| Top 19 | 60.55 | Top 53 | 61.47 | Top 88 | 63.30 | Top 123 | 60.55 | Top 158 | 62.39 |
| Top 20 | 60.55 | Top 54 | 64.22 | Top 89 | 64.22 | Top 124 | 60.55 | Top 159 | 62.39 |
| Top 21 | 58.72 | Top 55 | 62.39 | Top 90 | 63.30 | Top 126 | 57.80 | Top 160 | 61.47 |
| Top 22 | 61.47 | Top 56 | 62.39 | Top 91 | 63.30 | Top 127 | 58.72 | Top 161 | 62.39 |
| Top 23 | 57.80 | Top 57 | 61.47 | Top 92 | 63.30 | Top 128 | 56.88 | Top 162 | 61.47 |
| Top 24 | 55.96 | Top 58 | 60.55 | Top 93 | 66.97 | Top 129 | 55.96 | Top 163 | 62.39 |
| Top 25 | 55.96 | Top 59 | 61.47 | Top 94 | 66.06 | Top 130 | 55.96 | Top 164 | 62.39 |
| Top 26 | 56.88 | Top 60 | 62.39 | Top 95 | 64.22 | Top 131 | 57.80 | Top 165 | 62.39 |
| Top 27 | 60.55 | Top 62 | 60.55 | Top 96 | 65.14 | Top 132 | 57.80 | Top 166 | 62.39 |
| Top 28 | 59.63 | Top 63 | 61.47 | Top 97 | 63.30 | Top 133 | 57.80 | Top 167 | 62.39 |
| Top 29 | 57.80 | Top 64 | 61.47 | Top 98 | 64.22 | Top 134 | 57.80 | Top 168 | 63.30 |
| Top 30 | 57.80 | Top 65 | 60.55 | Top 100 | 61.47 | Top 135 | 60.55 | Top 169 | 63.30 |
| Top 31 | 57.80 | Top 66 | 58.72 | Top 101 | 62.39 | Top 136 | 63.30 | Top 170 | 62.39 |
| Top 32 | 57.80 | Top 67 | 58.72 | Top 102 | 64.22 | Top 137 | 62.39 |  |  |
| Top 33 | 56.88 | Top 68 | 60.55 | Top 103 | 62.39 | Top 138 | 62.39 |  |  |

**TABLE S5:** Models’ performance built on State4 in MDD dataset.

| Model | Accuracy | Model | Accuracy | Model | Accuracy | Model | Accuracy | Model | Accuracy |
| --- | --- | --- | --- | --- | --- | --- | --- | --- | --- |
| Baseline | 62.03 | Top 34 | 62.03 | Top 70 | 60.76 | Top 104 | 60.76 | Top 139 | 63.29 |
| Top 1 | 45.57 | Top 35 | 59.49 | Top 71 | 62.03 | Top 105 | 59.49 | Top 141 | 61.39 |
| Top 2 | 61.39 | Top 37 | 65.82 | Top 72 | 61.39 | Top 106 | 60.76 | Top 142 | 62.03 |
| Top 3 | 58.23 | Top 38 | 65.19 | Top 73 | 62.66 | Top 107 | 56.96 | Top 143 | 63.92 |
| Top 4 | 56.96 | Top 39 | 65.82 | Top 74 | 62.03 | Top 108 | 56.33 | Top 144 | 64.56 |
| Top 5 | 56.33 | Top 40 | 67.09 | Top 75 | 62.66 | Top 109 | 55.06 | Top 145 | 63.92 |
| Top 6 | 64.56 | Top 41 | 64.56 | Top 76 | 59.49 | Top 110 | 55.06 | Top 146 | 65.19 |
| Top 7 | 65.19 | Top 42 | 62.66 | Top 77 | 60.76 | Top 111 | 56.33 | Top 147 | 64.56 |
| Top 8 | 64.56 | Top 43 | 63.92 | Top 78 | 62.66 | Top 112 | 58.23 | Top 148 | 62.66 |
| Top 9 | 58.86 | Top 44 | 63.29 | Top 79 | 60.13 | Top 113 | 58.86 | Top 149 | 60.13 |
| Top 10 | 67.72 | Top 45 | 62.03 | Top 80 | 60.76 | Top 114 | 58.23 | Top 150 | 60.76 |
| Top 11 | 60.13 | Top 46 | 60.76 | Top 81 | 58.86 | Top 115 | 62.03 | Top 151 | 60.13 |
| Top 12 | 61.39 | Top 47 | 63.29 | Top 82 | 58.86 | Top 116 | 62.66 | Top 152 | 59.49 |
| Top 13 | 63.29 | Top 48 | 60.76 | Top 83 | 60.76 | Top 117 | 62.03 | Top 153 | 58.23 |
| Top 14 | 66.46 | Top 49 | 61.39 | Top 84 | 60.76 | Top 118 | 61.39 | Top 154 | 59.49 |
| Top 15 | 62.66 | Top 50 | 62.66 | Top 85 | 61.39 | Top 119 | 60.76 | Top 155 | 59.49 |
| Top 16 | 59.49 | Top 51 | 60.13 | Top 86 | 59.49 | Top 120 | 63.29 | Top 156 | 58.86 |
| Top 17 | 58.86 | Top 52 | 63.29 | Top 87 | 60.76 | Top 121 | 62.03 | Top 157 | 60.13 |
| Top 18 | 58.23 | Top 53 | 62.66 | Top 88 | 60.13 | Top 122 | 61.39 | Top 158 | 58.86 |
| Top 19 | 56.96 | Top 54 | 60.76 | Top 89 | 59.49 | Top 123 | 63.92 | Top 159 | 58.23 |
| Top 20 | 58.86 | Top 56 | 59.49 | Top 90 | 58.23 | Top 124 | 63.29 | Top 160 | 57.59 |
| Top 21 | 58.86 | Top 57 | 60.76 | Top 91 | 58.23 | Top 125 | 63.92 | Top 161 | 58.23 |
| Top 22 | 60.76 | Top 58 | 60.76 | Top 92 | 59.49 | Top 126 | 65.19 | Top 162 | 58.23 |
| Top 23 | 62.03 | Top 59 | 60.13 | Top 93 | 59.49 | Top 127 | 65.19 | Top 163 | 58.86 |
| Top 24 | 64.56 | Top 60 | 60.76 | Top 94 | 61.39 | Top 128 | 64.56 | Top 164 | 57.59 |
| Top 25 | 61.39 | Top 61 | 57.59 | Top 95 | 64.56 | Top 129 | 63.29 | Top 165 | 58.23 |
| Top 26 | 64.56 | Top 62 | 60.13 | Top 96 | 62.66 | Top 130 | 63.92 | Top 166 | 58.86 |
| Top 27 | 60.13 | Top 63 | 62.03 | Top 97 | 64.56 | Top 132 | 61.39 | Top 167 | 57.59 |
| Top 28 | 61.39 | Top 64 | 63.29 | Top 98 | 63.92 | Top 133 | 60.76 | Top 168 | 60.13 |
| Top 29 | 56.96 | Top 65 | 62.03 | Top 99 | 62.66 | Top 134 | 63.29 | Top 169 | 62.03 |
| Top 30 | 57.59 | Top 66 | 63.92 | Top 100 | 58.86 | Top 135 | 64.56 | Top 170 | 61.39 |
| Top 31 | 62.66 | Top 67 | 57.59 | Top 101 | 61.39 | Top 136 | 65.19 |  |  |
| Top 32 | 57.59 | Top 68 | 58.23 | Top 102 | 62.66 | Top 137 | 62.03 |  |  |
| Top 33 | 60.76 | Top 69 | 59.49 | Top 103 | 61.39 | Top 138 | 61.39 |  |  |

**TABLE S6:** Models’ performance built on State4 in BD dataset.

| Model | Accuracy | Model | Accuracy | Model | Accuracy | Model | Accuracy | Model | Accuracy |
| --- | --- | --- | --- | --- | --- | --- | --- | --- | --- |
| Baseline | 57.80 | Top 34 | 68.81 | Top 70 | 64.22 | Top 104 | 64.22 | Top 139 | 58.72 |
| Top 1 | 49.54 | Top 35 | 66.97 | Top 71 | 64.22 | Top 105 | 63.30 | Top 141 | 62.39 |
| Top 2 | 59.63 | Top 37 | 69.72 | Top 72 | 62.39 | Top 106 | 64.22 | Top 142 | 62.39 |
| Top 3 | 55.05 | Top 38 | 70.64 | Top 73 | 61.47 | Top 107 | 60.55 | Top 143 | 63.30 |
| Top 4 | 47.71 | Top 39 | 70.64 | Top 74 | 62.39 | Top 108 | 61.47 | Top 144 | 62.39 |
| Top 5 | 44.95 | Top 40 | 68.81 | Top 75 | 62.39 | Top 109 | 61.47 | Top 145 | 65.14 |
| Top 6 | 54.13 | Top 41 | 66.06 | Top 76 | 62.39 | Top 110 | 62.39 | Top 146 | 65.14 |
| Top 7 | 54.13 | Top 42 | 66.97 | Top 77 | 65.14 | Top 111 | 62.39 | Top 147 | 60.55 |
| Top 8 | 48.62 | Top 43 | 68.81 | Top 78 | 65.14 | Top 112 | 62.39 | Top 148 | 61.47 |
| Top 9 | 55.05 | Top 44 | 67.89 | Top 79 | 64.22 | Top 113 | 63.30 | Top 149 | 63.30 |
| Top 10 | 61.47 | Top 45 | 67.89 | Top 80 | 65.14 | Top 114 | 63.30 | Top 150 | 60.55 |
| Top 11 | 62.39 | Top 46 | 67.89 | Top 81 | 65.14 | Top 115 | 65.14 | Top 151 | 60.55 |
| Top 12 | 61.47 | Top 47 | 66.97 | Top 82 | 65.14 | Top 116 | 65.14 | Top 152 | 60.55 |
| Top 13 | 61.47 | Top 48 | 66.06 | Top 83 | 62.39 | Top 117 | 65.14 | Top 153 | 61.47 |
| Top 14 | 55.96 | Top 49 | 64.22 | Top 84 | 64.22 | Top 118 | 65.14 | Top 154 | 61.47 |
| Top 15 | 57.80 | Top 50 | 69.72 | Top 85 | 62.39 | Top 119 | 65.14 | Top 155 | 60.55 |
| Top 16 | 60.55 | Top 51 | 68.81 | Top 86 | 62.39 | Top 120 | 65.14 | Top 156 | 61.47 |
| Top 17 | 62.39 | Top 52 | 67.89 | Top 87 | 63.30 | Top 121 | 62.39 | Top 157 | 62.39 |
| Top 18 | 56.88 | Top 53 | 69.72 | Top 88 | 64.22 | Top 122 | 62.39 | Top 158 | 62.39 |
| Top 19 | 55.05 | Top 54 | 67.89 | Top 89 | 65.14 | Top 123 | 62.39 | Top 159 | 61.47 |
| Top 20 | 57.80 | Top 56 | 66.97 | Top 90 | 66.06 | Top 124 | 62.39 | Top 160 | 60.55 |
| Top 21 | 59.63 | Top 57 | 66.06 | Top 91 | 66.06 | Top 125 | 62.39 | Top 161 | 60.55 |
| Top 22 | 60.55 | Top 58 | 65.14 | Top 92 | 66.06 | Top 126 | 62.39 | Top 162 | 58.72 |
| Top 23 | 63.30 | Top 59 | 64.22 | Top 93 | 66.97 | Top 127 | 59.63 | Top 163 | 59.63 |
| Top 24 | 62.39 | Top 60 | 66.06 | Top 94 | 67.89 | Top 128 | 61.47 | Top 164 | 56.88 |
| Top 25 | 60.55 | Top 61 | 64.22 | Top 95 | 63.30 | Top 129 | 64.22 | Top 165 | 60.55 |
| Top 26 | 62.39 | Top 62 | 64.22 | Top 96 | 65.14 | Top 130 | 61.47 | Top 166 | 60.55 |
| Top 27 | 64.22 | Top 63 | 64.22 | Top 97 | 62.39 | Top 132 | 59.63 | Top 167 | 60.55 |
| Top 28 | 67.89 | Top 64 | 66.97 | Top 98 | 66.97 | Top 133 | 60.55 | Top 168 | 61.47 |
| Top 29 | 66.06 | Top 65 | 65.14 | Top 99 | 66.97 | Top 134 | 58.72 | Top 169 | 63.30 |
| Top 30 | 66.06 | Top 66 | 65.14 | Top 100 | 65.14 | Top 135 | 57.80 | Top 170 | 61.47 |
| Top 31 | 66.06 | Top 67 | 65.14 | Top 101 | 65.14 | Top 136 | 56.88 |  |  |
| Top 32 | 65.14 | Top 68 | 66.97 | Top 102 | 64.22 | Top 137 | 59.63 |  |  |
| Top 33 | 66.06 | Top 69 | 64.22 | Top 103 | 64.22 | Top 138 | 61.47 |  |  |

**TABLE S7:** Models’ performance built on State23 in MDD dataset.

| Model | Accuracy | Model | Accuracy | Model | Accuracy | Model | Accuracy | Model | Accuracy |
| --- | --- | --- | --- | --- | --- | --- | --- | --- | --- |
| Baseline | 62.03 | Top 33 | 62.66 | Top 67 | 54.43 | Top 102 | 56.33 | Top 139 | 61.39 |
| Top 1 | 60.76 | Top 34 | 65.19 | Top 68 | 54.43 | Top 103 | 58.23 | Top 140 | 59.49 |
| Top 2 | 56.33 | Top 35 | 63.29 | Top 69 | 53.80 | Top 104 | 57.59 | Top 141 | 58.86 |
| Top 3 | 66.46 | Top 36 | 60.76 | Top 70 | 62.03 | Top 105 | 56.96 | Top 142 | 60.76 |
| Top 4 | 69.62 | Top 37 | 64.56 | Top 73 | 59.49 | Top 106 | 59.49 | Top 144 | 58.86 |
| Top 5 | 56.96 | Top 38 | 61.39 | Top 74 | 58.86 | Top 107 | 58.86 | Top 145 | 59.49 |
| Top 6 | 61.39 | Top 39 | 59.49 | Top 75 | 58.23 | Top 108 | 60.13 | Top 146 | 60.76 |
| Top 7 | 59.49 | Top 40 | 60.76 | Top 76 | 55.70 | Top 109 | 59.49 | Top 147 | 60.76 |
| Top 8 | 62.03 | Top 41 | 59.49 | Top 77 | 57.59 | Top 111 | 58.86 | Top 148 | 57.59 |
| Top 9 | 60.13 | Top 42 | 59.49 | Top 78 | 59.49 | Top 112 | 60.13 | Top 149 | 60.76 |
| Top 10 | 65.82 | Top 43 | 62.66 | Top 79 | 60.13 | Top 113 | 58.23 | Top 150 | 58.23 |
| Top 11 | 59.49 | Top 44 | 63.29 | Top 80 | 58.23 | Top 114 | 56.96 | Top 151 | 56.33 |
| Top 12 | 65.19 | Top 45 | 63.29 | Top 81 | 56.96 | Top 115 | 61.39 | Top 152 | 53.80 |
| Top 13 | 62.03 | Top 46 | 65.19 | Top 82 | 57.59 | Top 116 | 61.39 | Top 153 | 53.16 |
| Top 14 | 63.29 | Top 47 | 66.46 | Top 83 | 55.70 | Top 117 | 60.13 | Top 154 | 54.43 |
| Top 15 | 64.56 | Top 48 | 67.72 | Top 84 | 59.49 | Top 119 | 58.23 | Top 155 | 54.43 |
| Top 16 | 62.66 | Top 49 | 65.19 | Top 85 | 56.96 | Top 120 | 59.49 | Top 156 | 55.70 |
| Top 17 | 60.76 | Top 50 | 65.82 | Top 86 | 57.59 | Top 121 | 58.23 | Top 157 | 56.33 |
| Top 18 | 57.59 | Top 51 | 66.46 | Top 87 | 56.33 | Top 122 | 58.23 | Top 158 | 57.59 |
| Top 19 | 60.76 | Top 52 | 65.19 | Top 88 | 55.06 | Top 123 | 57.59 | Top 159 | 55.70 |
| Top 20 | 61.39 | Top 53 | 65.82 | Top 89 | 57.59 | Top 124 | 60.76 | Top 160 | 55.70 |
| Top 21 | 62.66 | Top 54 | 64.56 | Top 90 | 57.59 | Top 125 | 60.13 | Top 161 | 57.59 |
| Top 22 | 63.92 | Top 55 | 62.66 | Top 91 | 58.23 | Top 126 | 60.13 | Top 162 | 58.23 |
| Top 23 | 63.92 | Top 56 | 60.13 | Top 92 | 57.59 | Top 127 | 56.96 | Top 163 | 57.59 |
| Top 24 | 60.76 | Top 57 | 58.86 | Top 93 | 60.13 | Top 128 | 56.96 | Top 164 | 55.06 |
| Top 25 | 68.35 | Top 58 | 61.39 | Top 94 | 58.86 | Top 129 | 56.33 | Top 165 | 57.59 |
| Top 26 | 58.23 | Top 59 | 56.96 | Top 95 | 58.23 | Top 130 | 58.86 | Top 166 | 58.23 |
| Top 27 | 67.09 | Top 60 | 61.39 | Top 96 | 58.86 | Top 131 | 58.86 | Top 167 | 58.23 |
| Top 28 | 63.29 | Top 61 | 59.49 | Top 97 | 56.96 | Top 132 | 58.86 | Top 168 | 58.23 |
| Top 29 | 63.29 | Top 62 | 58.86 | Top 98 | 60.13 | Top 133 | 60.13 | Top 169 | 58.86 |
| Top 30 | 67.09 | Top 63 | 57.59 | Top 99 | 59.49 | Top 134 | 59.49 | Top 170 | 58.86 |
| Top 31 | 67.72 | Top 65 | 56.96 | Top 100 | 57.59 | Top 135 | 58.86 |  |  |
| Top 32 | 66.46 | Top 66 | 56.96 | Top 101 | 57.59 | Top 138 | 60.13 |  |  |

**TABLE S8:** Models’ performance built on State23 in BD dataset.

| Model | Accuracy | Model | Accuracy | Model | Accuracy | Model | Accuracy | Model | Accuracy |
| --- | --- | --- | --- | --- | --- | --- | --- | --- | --- |
| Baseline | 57.80 | Top 33 | 64.22 | Top 67 | 61.47 | Top 102 | 64.22 | Top 139 | 65.14 |
| Top 1 | 57.80 | Top 34 | 60.55 | Top 68 | 59.63 | Top 103 | 64.22 | Top 140 | 65.14 |
| Top 2 | 66.06 | Top 35 | 60.55 | Top 69 | 62.39 | Top 104 | 64.22 | Top 141 | 66.06 |
| Top 3 | 64.22 | Top 36 | 63.30 | Top 70 | 65.14 | Top 105 | 64.22 | Top 142 | 65.14 |
| Top 4 | 64.22 | Top 37 | 65.14 | Top 73 | 64.22 | Top 106 | 64.22 | Top 144 | 64.22 |
| Top 5 | 62.39 | Top 38 | 66.06 | Top 74 | 64.22 | Top 107 | 65.14 | Top 145 | 63.30 |
| Top 6 | 57.80 | Top 39 | 62.39 | Top 75 | 65.14 | Top 108 | 61.47 | Top 146 | 62.39 |
| Top 7 | 62.39 | Top 40 | 64.22 | Top 76 | 65.14 | Top 109 | 65.14 | Top 147 | 62.39 |
| Top 8 | 63.30 | Top 41 | 60.55 | Top 77 | 66.06 | Top 111 | 64.22 | Top 148 | 61.47 |
| Top 9 | 64.22 | Top 42 | 63.30 | Top 78 | 67.89 | Top 112 | 59.63 | Top 149 | 61.47 |
| Top 10 | 63.30 | Top 43 | 64.22 | Top 79 | 67.89 | Top 113 | 59.63 | Top 150 | 62.39 |
| Top 11 | 64.22 | Top 44 | 63.30 | Top 80 | 67.89 | Top 114 | 59.63 | Top 151 | 65.14 |
| Top 12 | 65.14 | Top 45 | 63.30 | Top 81 | 67.89 | Top 115 | 62.39 | Top 152 | 64.22 |
| Top 13 | 62.39 | Top 46 | 60.55 | Top 82 | 66.06 | Top 116 | 60.55 | Top 153 | 66.06 |
| Top 14 | 60.55 | Top 47 | 62.39 | Top 83 | 67.89 | Top 117 | 58.72 | Top 154 | 61.47 |
| Top 15 | 62.39 | Top 48 | 65.14 | Top 84 | 68.81 | Top 119 | 60.55 | Top 155 | 62.39 |
| Top 16 | 61.47 | Top 49 | 61.47 | Top 85 | 65.14 | Top 120 | 60.55 | Top 156 | 66.97 |
| Top 17 | 63.30 | Top 50 | 63.30 | Top 86 | 63.30 | Top 121 | 60.55 | Top 157 | 66.06 |
| Top 18 | 63.30 | Top 51 | 60.55 | Top 87 | 64.22 | Top 122 | 64.22 | Top 158 | 63.30 |
| Top 19 | 61.47 | Top 52 | 61.47 | Top 88 | 65.14 | Top 123 | 63.30 | Top 159 | 64.22 |
| Top 20 | 60.55 | Top 53 | 60.55 | Top 89 | 64.22 | Top 124 | 65.14 | Top 160 | 63.30 |
| Top 21 | 63.30 | Top 54 | 59.63 | Top 90 | 65.14 | Top 125 | 66.06 | Top 161 | 64.22 |
| Top 22 | 63.30 | Top 55 | 63.30 | Top 91 | 65.14 | Top 126 | 64.22 | Top 162 | 62.39 |
| Top 23 | 65.14 | Top 56 | 63.30 | Top 92 | 64.22 | Top 127 | 62.39 | Top 163 | 62.39 |
| Top 24 | 66.97 | Top 57 | 63.30 | Top 93 | 65.14 | Top 128 | 63.30 | Top 164 | 62.39 |
| Top 25 | 64.22 | Top 58 | 63.30 | Top 94 | 64.22 | Top 129 | 63.30 | Top 165 | 65.14 |
| Top 26 | 63.30 | Top 59 | 62.39 | Top 95 | 65.14 | Top 130 | 64.22 | Top 166 | 63.30 |
| Top 27 | 58.72 | Top 60 | 62.39 | Top 96 | 65.14 | Top 131 | 62.39 | Top 167 | 61.47 |
| Top 28 | 61.47 | Top 61 | 61.47 | Top 97 | 64.22 | Top 132 | 61.47 | Top 168 | 61.47 |
| Top 29 | 62.39 | Top 62 | 62.39 | Top 98 | 62.39 | Top 133 | 65.14 | Top 169 | 60.55 |
| Top 30 | 57.80 | Top 63 | 63.30 | Top 99 | 62.39 | Top 134 | 64.22 | Top 170 | 63.30 |
| Top 31 | 60.55 | Top 65 | 63.30 | Top 100 | 62.39 | Top 135 | 63.30 |  |  |
| Top 32 | 60.55 | Top 66 | 60.55 | Top 101 | 64.22 | Top 138 | 65.14 |  |  |

**TABLE S9:**
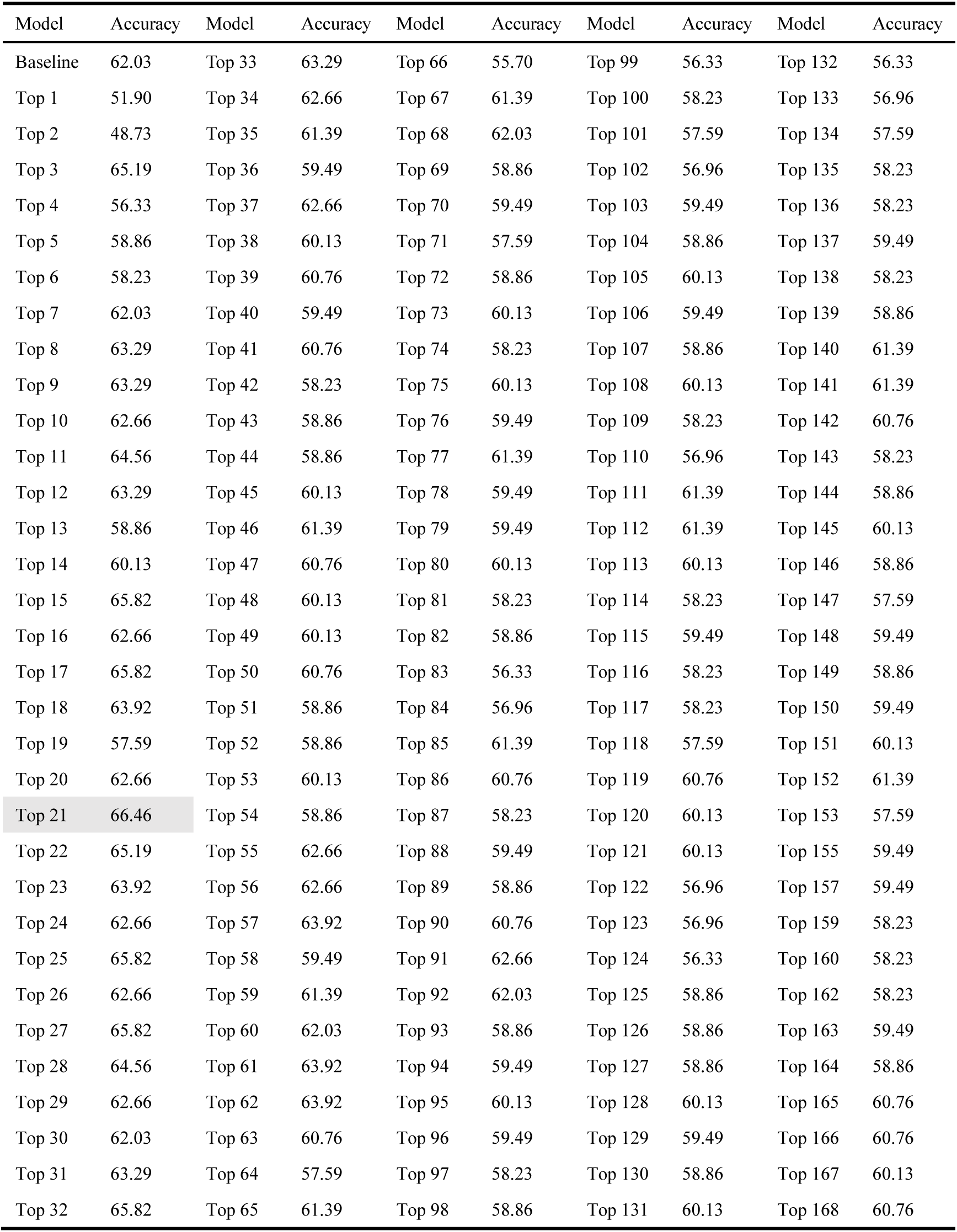
Models’ performance built on State234 in MDD dataset.

**TABLE S10:**
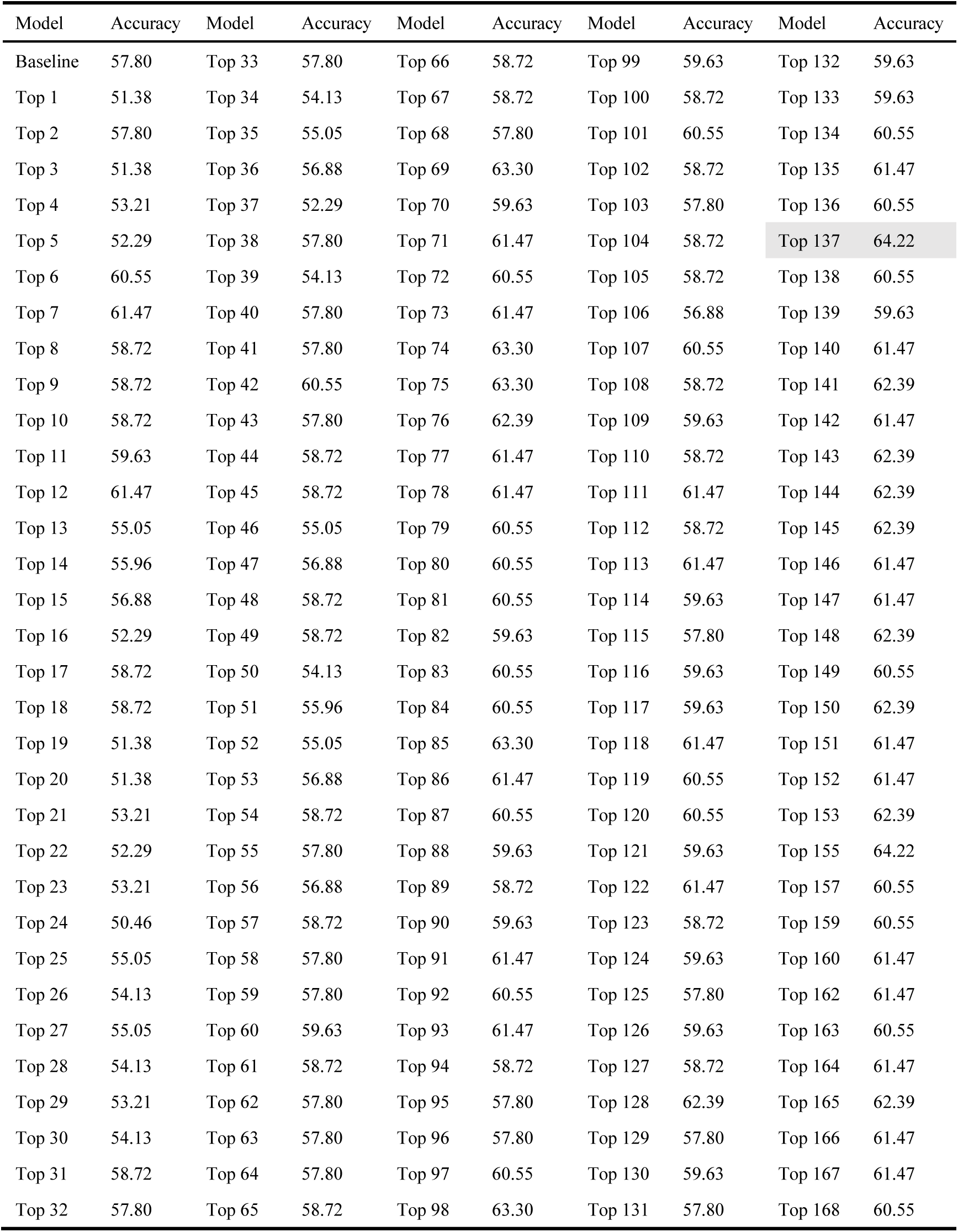
Models’ performance built on State234 in BD dataset.

| Model | Accuracy | Model | Accuracy | Model | Accuracy | Model | Accuracy | Model | Accuracy |
| --- | --- | --- | --- | --- | --- | --- | --- | --- | --- |
| Baseline | 57.80 | Top 33 | 57.80 | Top 66 | 58.72 | Top 99 | 59.63 | Top 132 | 59.63 |
| Top 1 | 51.38 | Top 34 | 54.13 | Top 67 | 58.72 | Top 100 | 58.72 | Top 133 | 59.63 |
| Top 2 | 57.80 | Top 35 | 55.05 | Top 68 | 57.80 | Top 101 | 60.55 | Top 134 | 60.55 |
| Top 3 | 51.38 | Top 36 | 56.88 | Top 69 | 63.30 | Top 102 | 58.72 | Top 135 | 61.47 |
| Top 4 | 53.21 | Top 37 | 52.29 | Top 70 | 59.63 | Top 103 | 57.80 | Top 136 | 60.55 |
| Top 5 | 52.29 | Top 38 | 57.80 | Top 71 | 61.47 | Top 104 | 58.72 | Top 137 | 64.22 |
| Top 6 | 60.55 | Top 39 | 54.13 | Top 72 | 60.55 | Top 105 | 58.72 | Top 138 | 60.55 |
| Top 7 | 61.47 | Top 40 | 57.80 | Top 73 | 61.47 | Top 106 | 56.88 | Top 139 | 59.63 |
| Top 8 | 58.72 | Top 41 | 57.80 | Top 74 | 63.30 | Top 107 | 60.55 | Top 140 | 61.47 |
| Top 9 | 58.72 | Top 42 | 60.55 | Top 75 | 63.30 | Top 108 | 58.72 | Top 141 | 62.39 |
| Top 10 | 58.72 | Top 43 | 57.80 | Top 76 | 62.39 | Top 109 | 59.63 | Top 142 | 61.47 |
| Top 11 | 59.63 | Top 44 | 58.72 | Top 77 | 61.47 | Top 110 | 58.72 | Top 143 | 62.39 |
| Top 12 | 61.47 | Top 45 | 58.72 | Top 78 | 61.47 | Top 111 | 61.47 | Top 144 | 62.39 |
| Top 13 | 55.05 | Top 46 | 55.05 | Top 79 | 60.55 | Top 112 | 58.72 | Top 145 | 62.39 |
| Top 14 | 55.96 | Top 47 | 56.88 | Top 80 | 60.55 | Top 113 | 61.47 | Top 146 | 61.47 |
| Top 15 | 56.88 | Top 48 | 58.72 | Top 81 | 60.55 | Top 114 | 59.63 | Top 147 | 61.47 |
| Top 16 | 52.29 | Top 49 | 58.72 | Top 82 | 59.63 | Top 115 | 57.80 | Top 148 | 62.39 |
| Top 17 | 58.72 | Top 50 | 54.13 | Top 83 | 60.55 | Top 116 | 59.63 | Top 149 | 60.55 |
| Top 18 | 58.72 | Top 51 | 55.96 | Top 84 | 60.55 | Top 117 | 59.63 | Top 150 | 62.39 |
| Top 19 | 51.38 | Top 52 | 55.05 | Top 85 | 63.30 | Top 118 | 61.47 | Top 151 | 61.47 |
| Top 20 | 51.38 | Top 53 | 56.88 | Top 86 | 61.47 | Top 119 | 60.55 | Top 152 | 61.47 |
| Top 21 | 53.21 | Top 54 | 58.72 | Top 87 | 60.55 | Top 120 | 60.55 | Top 153 | 62.39 |
| Top 22 | 52.29 | Top 55 | 57.80 | Top 88 | 59.63 | Top 121 | 59.63 | Top 155 | 64.22 |
| Top 23 | 53.21 | Top 56 | 56.88 | Top 89 | 58.72 | Top 122 | 61.47 | Top 157 | 60.55 |
| Top 24 | 50.46 | Top 57 | 58.72 | Top 90 | 59.63 | Top 123 | 58.72 | Top 159 | 60.55 |
| Top 25 | 55.05 | Top 58 | 57.80 | Top 91 | 61.47 | Top 124 | 59.63 | Top 160 | 61.47 |
| Top 26 | 54.13 | Top 59 | 57.80 | Top 92 | 60.55 | Top 125 | 57.80 | Top 162 | 61.47 |
| Top 27 | 55.05 | Top 60 | 59.63 | Top 93 | 61.47 | Top 126 | 59.63 | Top 163 | 60.55 |
| Top 28 | 54.13 | Top 61 | 58.72 | Top 94 | 58.72 | Top 127 | 58.72 | Top 164 | 61.47 |
| Top 29 | 53.21 | Top 62 | 57.80 | Top 95 | 57.80 | Top 128 | 62.39 | Top 165 | 62.39 |
| Top 30 | 54.13 | Top 63 | 57.80 | Top 96 | 57.80 | Top 129 | 57.80 | Top 166 | 61.47 |
| Top 31 | 58.72 | Top 64 | 57.80 | Top 97 | 60.55 | Top 130 | 59.63 | Top 167 | 61.47 |
| Top 32 | 57.80 | Top 65 | 58.72 | Top 98 | 63.30 | Top 131 | 57.80 | Top 168 | 60.55 |

### Self-acquired healthy dataset

#### Stimuli

This study used movies that can strongly induce subjects’ emotional experiences as emotional stimulus materials. The emotional stimulus materials were 12 different 10-minute naturalistic episode candidates with strong and reliable inducing effects of happy and sad emotional experiences, which included 6 happy clips (happy emotions) and 6 sad clips (sad emotions). There was no content overlap between the selected episodes. The 6 happy episodes are from the movies of “Mr. Popper’s Penguins”, “Ted”, “The Onion Movie”, “Liar Liar”, “A Thousand Words”, and “Absolutely Anything”, and the 6 sad episodes are from “Miracle In Cell No.7”, “Prayers For Bobby”, “The Classic”, “Grave Of The Fireflies”, “Only The Brave”, and “The Last Train”.

#### Experimental paradigm

The experimental paradigm is shown in Supplementary Fig S2. For each subject, the entire experiment consisted of a total of 6 trials (corresponding to 6 episodes). The 6 episodes were randomly selected from 12 different 10-minute episode candidates, including 3 happy and 3 sad stimuli. The sequence of the presentation was randomized to balance the sequential effect. Similar to previous studies, episodes were presented without sound. Each episode was subtitled to facilitate understanding of the content. Each trial consisted of a 30-second baseline (subjects watched a white cross displayed in the center of a black screen), a 10-minute movie playback (subjects were fully engaged in passively watching the episode being played), and subjective feedback (subjects rated their emotional experience evoked by the presented episodes using a 5-point happy-sad scale). Throughout the procedure, subjects were asked to remain stationary. The entire experimental paradigm was assessed by E-Prime 3.0 software.

#### Data acquisition

Brain images were obtained using a 64-channel head coil on a 3-Tesla Siemens Prisma MRI scanner. High-resolution T1-weighted structural images were acquired using a magnetization-prepared rapid acquisition gradient echo (MPRAGE) sequence with voxel resolution=1×1×1 mm^3^, repetition time (TR)=2300 ms, echo time (TE)=2.26 ms, field of view (FOV) =256×232 mm^2^, flip angle (FA)=8°. The functional images were recorded using a single gradient echo-planar imaging (EPI) sequence, with TR=1000ms, TE=30ms, FOV=192×192mm^2^, FA=90°, with a high spatial resolution of 2×2×2 mm^3^. Each volume of EPI functional images consisted of 65 slices, the total volume is 630 including 600 volumes during movie watching and 30 volumes during cross gazing. During scanning, all subjects were instructed to remain awake, keep their eyes open, and be in full engagement with the presented episodes. episodes were counter-projected on a screen and viewed through a mirror mounted on a head coil.

### Public MDD dataset

#### Data acquisition

Brain images were obtained using a 12-channel head coil on a 3-Tesla Siemens Magnetom Verio.Dot MRI scanner. The functional images were recorded using a single gradient echo-planar imaging (EPI) sequence, with TR=2500ms, TE=30ms, FOV=212×212mm^2^, FA=80 °, with a spatial resolution of 3.3×3.3×4mm^3^. Each volume of EPI functional images consisted of 40 slices, total volume=240+4 (dummy). During scanning (10 minutes and 10 seconds dummy period), all participants were instructed to remain awake, look at the fixation point and not think about anything in particular.

**Fig S2:**
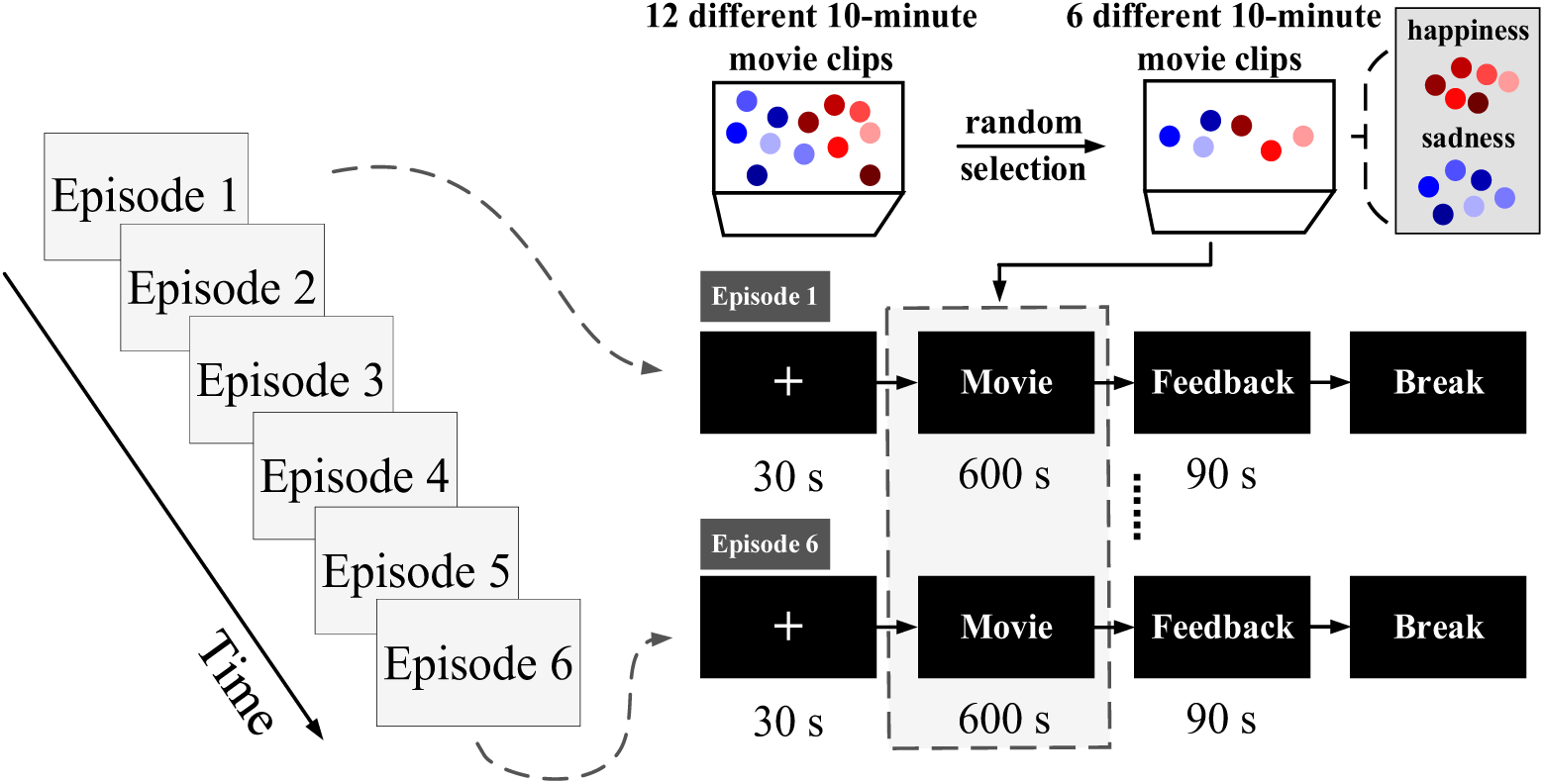
The experimental paradigm. For each subject, the experiment included 6 movie clips (episodes) presented in a randomized order. For each subject, 6 movie clips, including 3 happy movie clips and 3 sad movie clips, were randomly selected from a total of 12 different 10-minute movie clips.

### Self-acquired BD dataset

#### Data acquisition

The imaging data were performed on a Siemens 3T Trio scanner with a 12-channel head coil. Resting-state fMRI data were acquired using a standard gradient-echo EPI sequence with 31 oblique slices, TR=2000ms, TE=30ms, FOV=240×240mm^2^, FA= 90°, voxel size=3×3×5mm^3^, total volume=246. During the whole scan, all participants were requested to keep their eyes open and stay awake.

## Notes

### Competing Interest Statement

The authors have declared no competing interest.

## References

Abrol, A., Damaraju, E., Miller, R.L., et al., Replicability of time-varying connectivity patterns in large resting state fMRI samples.[J] Neuroimage, 2017. 163: p. 160-176.

Banks, S.J., Eddy, K.T., Angstadt, M., et al., Amygdala–frontal connectivity during emotion regulation.[J] Social cognitive and affective neuroscience, 2007. 2(4): p. 303-312.

Berridge, K.C., and Kringelbach, M.L., Building a neuroscience of pleasure and well-being.[J] Psychology of Well-Being: Theory, Research and Practice, 2011. 1: p. 1-26.

Berridge, K.C., and Kringelbach, M.L., Neuroscience of affect: brain mechanisms of pleasure and displeasure.[J] Current opinion in neurobiology, 2013. 23(3): p. 294–303.

Bijsterbosch, J., Smith, S.M., and Beckmann, C., An introduction to resting state fMRI functional connectivity[M]. Oxford University Press. 2017.

Bondi, E., Maggioni, E., Brambilla, P., et al., A systematic review on the potential use of machine learning to classify major depressive disorder from healthy controls using resting state fMRI measures.[J] Neuroscience & Biobehavioral Reviews, 2023. 144: p. 104972.

Bore, M.C., Liu, X., Gan, X., et al., Distinct neurofunctional alterations during motivational and hedonic processing of natural and monetary rewards in depression–a neuroimaging meta-analysis.[J] Psychological Medicine, 2024. 54(4): p. 639–651.

Bore, M.C., Liu, X., Huang, X., et al., Common and separable neural alterations in adult and adolescent depression–evidence from neuroimaging meta-analyses.[J] Neuroscience & Biobehavioral Reviews, 2024: p. 105835.

Buckner, R.L., Andrews-Hanna, J.R., and Schacter, D.L., The brain’s default network: anatomy, function, and relevance to disease.[J] Annals of the new York Academy of Sciences, 2008. 1124(1): p. 1–38.

Canario, E., Chen, D., and Biswal, B., A review of resting-state fMRI and its use to examine psychiatric disorders.[J] Psychoradiology, 2021. 1(1): p. 42–53.

Čeko, M., Kragel, P.A., Woo, C.-W., et al., Common and stimulus-type-specific brain representations of negative affect.[J] Nature neuroscience, 2022. 25(6): p. 760-770.

Chen, Y., Liu, C., Xin, F., et al., Opposing and emotion-specific associations between frontal activation with depression and anxiety symptoms during facial emotion processing in generalized anxiety and depression.[J] Progress in Neuro-Psychopharmacology and Biological Psychiatry, 2023. 123: p. 110716.

Cole, M.W., Yarkoni, T., Repovš, G., et al., Global connectivity of prefrontal cortex predicts cognitive control and intelligence.[J] Journal of Neuroscience, 2012. 32(26): p. 8988–8999.

Collaborators, G.M.D., Global, regional, and national burden of 12 mental disorders in 204 countries and territories, 1990–2019: a systematic analysis for the Global Burden of Disease Study 2019.[J] The Lancet Psychiatry, 2022. 9(2): p. 137–150.

Costello, E.J., Pine, D.S., Hammen, C., et al., Development and natural history of mood disorders.[J] Biological psychiatry, 2002. 52(6): p. 529–542.

Cromby, J., and Willis, M.E., Affect—or feeling (after Leys).[J] Theory & Psychology, 2016. 26(4): p. 476-495.

Dai, P., Zhou, X., Xiong, T., et al., Altered effective connectivity among the cerebellum and cerebrum in patients with major depressive disorder using multisite resting-state fMRI.[J] The Cerebellum, 2023. 22(5): p. 781-789.

Dai, P., Zhou, Y., Shi, Y., et al., Classification of MDD using a Transformer classifier with large-scale multisite resting-state fMRI data.[J] Human brain mapping, 2024. 45(1): p. e26542.

Dodd, A., Lockwood, E., Mansell, W., et al., Emotion regulation strategies in bipolar disorder: A systematic and critical review.[J] Journal of affective disorders, 2019. 246: p. 262–284.

Du, Y., Fryer, S.L., Fu, Z., et al., Dynamic functional connectivity impairments in early schizophrenia and clinical high-risk for psychosis.[J] Neuroimage, 2018. 180: p. 632–645.

Eickhoff, S.B., Milham, M., and Vanderwal, T., Towards clinical applications of movie fMRI.[J] NeuroImage, 2020. 217: p. 116860.

Etkin, A., A reckoning and research agenda for neuroimaging in psychiatry.[J] American Journal of Psychiatry, 2019. 176(7): p. 507–511.

Fekadu, N., Shibeshi, W., and Engidawork, E., Major depressive disorder: pathophysiology and clinical management.[J] J Depress Anxiety, 2017. 6(1): p. 255–257.

Firouzi, M., Kazemi, K., Ahmadi, M., et al., Enhanced ADHD classification through deep learning and dynamic resting state fMRI analysis.[J] Scientific Reports, 2024. 14(1): p. 24473.

Friston, K.J., Statistical parametric mapping.[J] Neuroscience databases: a practical guide, 2003: p. 237–250.

Gallo, S., El-Gazzar, A., Zhutovsky, P., et al., Functional connectivity signatures of major depressive disorder: machine learning analysis of two multicenter neuroimaging studies.[J] Molecular Psychiatry, 2023. 28(7): p. 3013–3022.

Gan, X., Zhou, X., Li, J., et al., Common and distinct neurofunctional representations of core and social disgust in the brain: Coordinate-based and network meta-analyses.[J] Neuroscience & Biobehavioral Reviews, 2022. 135: p. 104553.

Guo, S., Zhao, W., Tao, H., et al., The instability of functional connectivity in patients with schizophrenia and their siblings: a dynamic connectivity study.[J] Schizophrenia Research, 2018. 195: p. 183-189.

Hasson, U., Nir, Y., Levy, I., et al., Intersubject synchronization of cortical activity during natural vision.[J] science, 2004. 303(5664): p. 1634-1640.

Ho, C.-Y., and Berridge, K.C., An orexin hotspot in ventral pallidum amplifies hedonic ‘liking’for sweetness.[J] Neuropsychopharmacology, 2013. 38(9): p. 1655–1664.

Hornak, J., Bramham, J., Rolls, E.T., et al., Changes in emotion after circumscribed surgical lesions of the orbitofrontal and cingulate cortices.[J] Brain, 2003. 126(7): p. 1691–1712.

Houenou, J., Frommberger, J., Carde, S., et al., Neuroimaging-based markers of bipolar disorder: evidence from two meta-analyses.[J] Journal of affective disorders, 2011. 132(3): p. 344–355.

Huskey, R., Craighead, B., Miller, M.B., et al., Does intrinsic reward motivate cognitive control? A naturalistic-fMRI study based on the synchronization theory of flow.[J] Cognitive, Affective, & Behavioral Neuroscience, 2018. 18: p. 902-924.

Insel, T., Cuthbert, B., Garvey, M., et al.,2010. Research domain criteria (RDoC): toward a new classification framework for research on mental disorders[J]. Journal. 167, 748–751.

Jaworska, N., Yang, X.-R., Knott, V., et al., A review of fMRI studies during visual emotive processing in major depressive disorder.[J] The World Journal of Biological Psychiatry, 2015. 16(7): p. 448–471.

Jiang, X., Ma, X., Geng, Y., et al., Intrinsic, dynamic and effective connectivity among large-scale brain networks modulated by oxytocin.[J] Neuroimage, 2021. 227: p. 117668.

Jie, N.-F., Osuch, E.A., Zhu, M.-H., et al., Discriminating bipolar disorder from major depression using whole-brain functional connectivity: a feature selection analysis with SVM-FoBa algorithm.[J] Journal of Signal Processing Systems, 2018. 90: p. 259–271.

Johnstone, T., Van Reekum, C.M., Urry, H.L., et al., Failure to regulate: counterproductive recruitment of top-down prefrontal-subcortical circuitry in major depression.[J] Journal of Neuroscience, 2007. 27(33): p. 8877–8884.

Kessler, R., Berglund, P., and Demler, O., Mood disorders: Bipolar and major depressive disorders.[J] Jama, 2003. 289(23): p. 3095–105.

Lawrence, N.S., Williams, A.M., Surguladze, S., et al., Subcortical and ventral prefrontal cortical neural responses to facial expressions distinguish patients with bipolar disorder and major depression.[J] Biological psychiatry, 2004. 55(6): p. 578-587.

Li, G.-z., Liu, P.-h., Zhang, A.-x., et al., A resting state fMRI study of major depressive disorder with and without anxiety.[J] Psychiatry research, 2022. 315: p. 114697.

Li, J., Chen, J., Kong, W., et al., Abnormal core functional connectivity on the pathology of MDD and antidepressant treatment: A systematic review.[J] Journal of affective disorders, 2022. 296: p. 622–634.

Liu, C., Dai, J., Chen, Y., et al., Disorder-and emotional context-specific neurofunctional alterations during inhibitory control in generalized anxiety and major depressive disorder.[J] NeuroImage: Clinical, 2021. 30: p. 102661.

Liu, M., Wang, Y., Zhang, A., et al., Altered dynamic functional connectivity across mood states in bipolar disorder.[J] Brain Research, 2021. 1750: p. 147143.

Liu, Q., Zhou, B., Zhang, X., et al., Abnormal multi-layered dynamic cortico-subcortical functional connectivity in major depressive disorder and generalized anxiety disorder.[J] Journal of Psychiatric Research, 2023. 167: p. 23–31.

Liu, X., Jiao, G., Zhou, F., et al., A neural signature for the subjective experience of threat anticipation under uncertainty.[J] Nature Communications, 2024. 15(1): p. 1544.

Lu, F., Chen, Y., Cui, Q., et al., Shared and distinct patterns of dynamic functional connectivity variability of thalamo-cortical circuit in bipolar depression and major depressive disorder.[J] Cerebral Cortex, 2023. 33(11): p. 6681-6692.

Lu, F., Cui, Q., He, Z., et al., Prefrontal-limbic-striatum dysconnectivity associated with negative emotional endophenotypes in bipolar disorder during depressive episodes.[J] Journal of affective disorders, 2021. 295: p. 422–430.

Luppi, A.I., Mediano, P.A., Rosas, F.E., et al., A synergistic core for human brain evolution and cognition.[J] Nature Neuroscience, 2022. 25(6): p. 771-782.

Luppi, A.I., and Stamatakis, E.A., Combining network topology and information theory to construct representative brain networks.[J] Network Neuroscience, 2021. 5(1): p. 96–124.

Mahler, S.V., Smith, K.S., and Berridge, K.C., Endocannabinoid hedonic hotspot for sensory pleasure: anandamide in nucleus accumbens shell enhances ‘liking’of a sweet reward.[J] Neuropsychopharmacology, 2007. 32(11): p. 2267–2278.

Malhi, G.S., Bell, E., Bassett, D., et al., The 2020 Royal Australian and New Zealand College of Psychiatrists clinical practice guidelines for mood disorders.[J] Australian & New Zealand Journal of Psychiatry, 2021. 55(1): p. 7-117.

Matsui, T., and Yamashita, K.-i., Static and dynamic functional connectivity alterations in Alzheimer’s disease and neuropsychiatric diseases.[J] Brain Connectivity, 2023. 13(5): p. 307–314.

Meer, J.N.v.d., Breakspear, M., Chang, L.J., et al., Movie viewing elicits rich and reliable brain state dynamics.[J] Nature communications, 2020. 11(1): p. 5004.

Moran, J.M., Macrae, C.N., Heatherton, T.F., et al., Neuroanatomical evidence for distinct cognitive and affective components of self.[J] Journal of cognitive neuroscience, 2006. 18(9): p. 1586–1594.

Mousavian, M., Chen, J., Traylor, Z., et al., Depression detection from sMRI and rs-fMRI images using machine learning.[J] Journal of Intelligent Information Systems, 2021. 57: p. 395–418.

Noah, J.A., Ono, Y., Nomoto, Y., et al., fMRI validation of fNIRS measurements during a naturalistic task.[J] JoVE (Journal of Visualized Experiments), 2015(100): p. e52116.

Ochsner, K.N., and Gross, J.J., The cognitive control of emotion.[J] Trends in cognitive sciences, 2005. 9(5): p. 242-249.

Öngür, D., Lundy, M., Greenhouse, I., et al., Default mode network abnormalities in bipolar disorder and schizophrenia.[J] Psychiatry Research: Neuroimaging, 2010. 183(1): p. 59–68.

Parkes, L., Satterthwaite, T.D., and Bassett, D.S., Towards precise resting-state fMRI biomarkers in psychiatry: synthesizing developments in transdiagnostic research, dimensional models of psychopathology, and normative neurodevelopment.[J] Current Opinion in Neurobiology, 2020. 65: p. 120–128.

Peng, X., Wu, X., Gong, R., et al., Sub-regional anterior cingulate cortex functional connectivity revealed default network subsystem dysfunction in patients with major depressive disorder.[J] Psychological Medicine, 2021. 51(10): p. 1687-1695.

Phillips, M.L., Travis, M.J., Fagiolini, A., et al., Medication effects in neuroimaging studies of bipolar disorder.[J] American Journal of Psychiatry, 2008. 165(3): p. 313–320.

Pilmeyer, J., Huijbers, W., Lamerichs, R., et al., Functional MRI in major depressive disorder: A review of findings, limitations, and future prospects.[J] Journal of Neuroimaging, 2022. 32(4): p. 582–595.

Pitsillou, E., Bresnehan, S.M., Kagarakis, E.A., et al., The cellular and molecular basis of major depressive disorder: towards a unified model for understanding clinical depression.[J] Molecular biology reports, 2020. 47: p. 753-770.

Pourtois, G., and Vuilleumier, P., Dynamics of emotional effects on spatial attention in the human visual cortex.[J] Progress in brain research, 2006. 156: p. 67–91.

Preti, M.G., Bolton, T.A., and Van De Ville, D., The dynamic functional connectome: State-of-the-art and perspectives.[J] Neuroimage, 2017. 160: p. 41–54.

Qiao, D., Zhang, A., Sun, N., et al., Altered static and dynamic functional connectivity of habenula associated with suicidal ideation in first-episode, drug-naïve patients with major depressive disorder.[J] Frontiers in Psychiatry, 2020. 11: p. 608197.

Ramirez-Mahaluf, J.P., Perramon, J., Otal, B., et al., Subgenual anterior cingulate cortex controls sadness-induced modulations of cognitive and emotional network hubs.[J] Scientific reports, 2018. 8(1): p. 8566.

Rolls, E.T., The orbitofrontal cortex.[J] Philosophical Transactions of the Royal Society of London. Series B: Biological Sciences, 1996. 351(1346): p. 1433-1444.

Saarimäki, H., Glerean, E., Smirnov, D., et al., Classification of emotion categories based on functional connectivity patterns of the human brain.[J] NeuroImage, 2022. 247: p. 118800.

Schaefer, A., Kong, R., Gordon, E.M., et al., Local-global parcellation of the human cerebral cortex from intrinsic functional connectivity MRI.[J] Cerebral cortex, 2018. 28(9): p. 3095-3114.

Siegle, G.J., Thompson, W., Carter, C.S., et al., Increased amygdala and decreased dorsolateral prefrontal BOLD responses in unipolar depression: related and independent features.[J] Biological psychiatry, 2007. 61(2): p. 198–209.

Sonkusare, S., Breakspear, M., and Guo, C., Naturalistic stimuli in neuroscience: critically acclaimed.[J] Trends in cognitive sciences, 2019. 23(8): p. 699–714.

Tanaka, S.C., Yamashita, A., Yahata, N., et al., A multi-site, multi-disorder resting-state magnetic resonance image database.[J] Scientific data, 2021. 8(1): p. 227.

Taschereau-Dumouchel, V., Michel, M., Lau, H., et al., Putting the “mental” back in “mental disorders”: a perspective from research on fear and anxiety.[J] Molecular Psychiatry, 2022. 27(3): p. 1322-1330.

Tian, Y., Margulies, D.S., Breakspear, M., et al., Topographic organization of the human subcortex unveiled with functional connectivity gradients.[J] Nature neuroscience, 2020. 23(11): p. 1421-1432.

Toh, W.L., Thomas, N., and Rossell, S.L., Auditory verbal hallucinations in bipolar disorder (BD) and major depressive disorder (MDD): A systematic review.[J] Journal of affective disorders, 2015. 184: p. 18–28.

Van Den Heuvel, M.P., and Pol, H.E.H., Exploring the brain network: a review on resting-state fMRI functional connectivity.[J] European neuropsychopharmacology, 2010. 20(8): p. 519–534.

Wang, J., Ren, Y., Hu, X., et al., Test–retest reliability of functional connectivity networks during naturalistic fMRI paradigms.[J] Human brain mapping, 2017. 38(4): p. 2226–2241.

Welton, T., Indja, B.E., Maller, J.J., et al., Replicable brain signatures of emotional bias and memory based on diffusion kurtosis imaging of white matter tracts.[J] Human brain mapping, 2020. 41(5): p. 1274-1285.

Wen, Y., Li, H., Huang, Y., et al., Dynamic network characteristics of adolescents with major depressive disorder: Attention network mediates the association between anhedonia and attentional deficit.[J] Human Brain Mapping, 2023. 44(17): p. 5749–5769.

Winter, N.R., Blanke, J., Leenings, R., et al., A systematic evaluation of machine learning–based biomarkers for major depressive disorder.[J] JAMA psychiatry, 2024. 81(4): p. 386-395.

Xu, S., Zhang, Z., Li, L., et al., Functional connectivity profiles of the default mode and visual networks reflect temporal accumulative effects of sustained naturalistic emotional experience.[J] NeuroImage, 2023. 269: p. 119941.

Xu, X., Dai, J., Chen, Y., et al., Intrinsic connectivity of the prefrontal cortex and striato-limbic system respectively differentiate major depressive from generalized anxiety disorder.[J] Neuropsychopharmacology, 2021. 46(4): p. 791-798.

Xu, X., Dai, J., Liu, C., et al., Common and disorder-specific neurofunctional markers of dysregulated empathic reactivity in major depression and generalized anxiety disorder.[J] Psychotherapy and psychosomatics, 2020. 89(2): p. 114-116.

Xu, X., Xin, F., Liu, C., et al., Disorder-and cognitive demand-specific neurofunctional alterations during social emotional working memory in generalized anxiety disorder and major depressive disorder.[J] Journal of Affective Disorders, 2022. 308: p. 98–105.

Yan, C.-G., Wang, X.-D., Zuo, X.-N., et al., DPABI: data processing & analysis for (resting-state) brain imaging.[J] Neuroinformatics, 2016. 14: p. 339–351.

Zhang, X., Liu, J., Yang, Y., et al., Test–retest reliability of dynamic functional connectivity in naturalistic paradigm functional magnetic resonance imaging.[J] Human brain mapping, 2022. 43(4): p. 1463–1476.

Zhao, C., Huang, W.-J., Feng, F., et al., Abnormal characterization of dynamic functional connectivity in Alzheimer’s disease.[J] Neural regeneration research, 2022. 17(9): p. 2014-2021.

Zhong, S., Chen, P., Lai, S., et al., Aberrant dynamic functional connectivity in corticostriatal circuitry in depressed bipolar II disorder with recent suicide attempt.[J] Journal of affective disorders, 2022. 319: p. 538–548.

Zhou, F., Zhang, R., Yao, S., et al., Capturing dynamic fear experiences in naturalistic contexts: An ecologically valid fMRI signature integrating brain activation and connectivity.[J] bioRxiv, 2023: p. 2023.08. 18.553808.

